# Reconstructing clone-resolved transcriptional programs from bulk tumor sequencing

**DOI:** 10.64898/2026.01.15.699695

**Authors:** Jiaying Lai, Yi Yang, Kathleen Noller, Yunzhou Liu, Archana Balan, Prathima B. Nagendra, Luciane T. Kagohara, Elana J. Fertig, Laura D. Wood, Rachel Karchin

## Abstract

Tumor clones acquire distinct transcriptional programs as they evolve, but bulk RNA-seq averages over clonal mixtures and obscures the lineage-specific biology that DNA-sequencing reveals. We present PICTographPlus, the first method to infer clone-resolved transcriptional programs by integrating bulk DNA-derived clonal phylogenies and proportions with bulk RNA-seq alone, without single-cell data. Benchmarked against experimentally measured ground truth, scDNA/scRNA co-profiled cells from a wellDR-seq cancer dataset, across 320 pseudo-bulk replicates spanning four tumor purities and four sample counts and evaluated under seven regularization models, PICTographPlus recovers clone-level expression at mean Pearson r ≥ 0.92 and localizes pathway gains and losses to correct evolutionary branches (median F1 0.31–0.40, well above a no-deconvolution baseline). Applied to multi-region NSCLC, pancreatic precursor lesions, and rapid-autopsy PDAC, it localizes metabolic reprogramming, precursor-to-invasive transitions, and organ-adapted metastatic states to specific clonal branches. PICTographPlus turns standard bulk assays into clone-resolved transcriptional maps, enabling retrospective analyses where single-cell profiling is impractical.

## Introduction

Tumor evolution and tumor transcription are typically studied with different assays. Bulk and single-cell DNA (scDNA) sequencing recover clonal architectures and evolutionary histories with increasing precision^1–3^. Single-cell and spatial RNA profiling reveal diverse transcriptional states and microenvironmental niches^4–6^. What has remained out of reach, and is the focus of this work, is reading clone-resolved transcriptional programs from the multi-region or longitudinal cohorts where bulk DNA and bulk RNA are jointly available but matched single-cell profiling is not. Doing so requires a quantitative model that decomposes a bulk RNA mixture into latent clone-specific profiles, leverages DNA-derived clone proportions and phylogeny as constraints, and projects the resulting transcriptional differences onto the evolutionary edges along which they were acquired.

Single-cell RNA sequencing (scRNA-seq) can in principle recover clone-level expression when somatic variants or copy-number alterations are linked to individual cells, but single-cell assays remain expensive, technically demanding, and sensitive to dissociation bias, limiting routine use in large retrospective cohorts^7^. Bulk RNA-seq is widely available but averages heterogeneous mixtures into a single expression profile per sample, obscuring lineage-specific programs.

Several approaches address related aspects, but none reconstruct clone-specific transcriptomes from bulk DNA and RNA while leveraging clonal phylogenies. Expression deconvolution methods such as CIBERSORTx and MuSiC separate broad cell types without modeling genetically defined clones^8,9^. Methods that integrate transcriptomes with DNA-defined clones --clonealign and cardelino, which assign single cells to clones, and Canopy2, PhylEx, and TUSV-int, which jointly infer clone phylogenies and clone-level expression, all require single-cell RNA ^10–14^. Clonal inference frameworks such as PhyloWGS and Canopy reconstruct trees from DNA alone, without recovering transcriptional programs^2,3,15,16^.

Here, we introduce PICTographPlus, a computational framework that integrates bulk DNA-based clonal reconstruction with bulk RNA-seq to infer clone-specific transcriptional programs without requiring single-cell data. PICTographPlus takes as input mutation and copy-number calls from multi-region tumor samples, clusters alterations into subclonal groups, and reconstructs an evolutionary tree using a 1-Dollo model^17^. It then links this clonal structure to bulk expression through a regularized mixture model that represents observed sample-level expression as a clone-weighted combination of latent clone-specific profiles. The clonal phylogeny serves as the scaffold onto which recovered clone-specific expression differences are mapped, enabling localization of gene- and pathway-level changes to specific evolutionary edges. To our knowledge, PICTographPlus is the first method to jointly leverage DNA-inferred clonal architectures and bulk RNA-seq to recover clone-resolved transcriptional programs and map them onto evolutionary branches from standard bulk assays.

We benchmark PICTographPlus using co-profiled scDNA/scRNA data from a patient of the breast cancer cohort profiled by wellDR-seq^18^, which provides jointly measured copy-number-defined clones and their transcriptomes for every cell. We extend the resulting four-clone tree with synthetic clones carrying known Hallmark gene set perturbations to create a deep-tree benchmark for systematic stress testing. To demonstrate the method on real bulk multi-region data, we then present three illustrative single-patient case studies, each chosen to span a distinct disease context: a TRACERx non-small cell lung cancer (NSCLC) case (primary-to-therapy-associated metastasis), a pancreatic precursor lesion from the NCI Cancer Moonshot Precancer Atlas Pilot (precursor-to-invasive progression), and a rapid-autopsy PDAC case from the PanCuRx Translational Research Initiative (multi-organ metastatic divergence)^19–23^. In each of these single-patient case studies, PICTographPlus recovered clone-specific gene set programs and linked DNA-defined evolutionary events to their transcriptional consequences, illustrating how the method can resolve precursor, primary, metastatic, and therapy-associated transitions from bulk DNA and RNA data alone. We emphasize that these are individual case applications intended to demonstrate the method on real data; quantitative performance is established by the 320-replicate benchmark above, and recurrence of the specific biological programs identified here will require future cohort-scale studies. More broadly, PICTographPlus demonstrates how clonal phylogenies can serve as a scaffold for integrating genomic evolution with transcriptional reprogramming. Because the model requires only a mixture of clone-specific profiles with known mixing proportions, the same formulation extends in principle to other mixed measurements, including single-cell pseudo-bulks and spatial transcriptomic spots, a direction we leave to future work.

## Results

### PICTographPlus framework

PICTographPlus performs joint inference across two genomic data modalities that have historically been analyzed in isolation. From bulk DNA, it derives a clonal phylogeny and per-sample clone proportions using a Bayesian generative model with Markov chain Monte Carlo sampling, followed by a modified Gabow-Myers tree enumeration (Methods; Note S1)^3,24,25^.

From bulk RNA-seq, it observes a clone-weighted mixture of latent clone-specific expression profiles. The framework couples these by treating the DNA-derived clone proportion matrix as a fixed constraint on the RNA mixture model, allowing the clone-by-gene expression matrix to be inferred as the solution to a regularized least-squares problem (Figure 1). Extending the PICTograph framework with this RNA-coupling step is what enables clone-resolved transcriptional inference from standard bulk assays.

**Figure 1.**
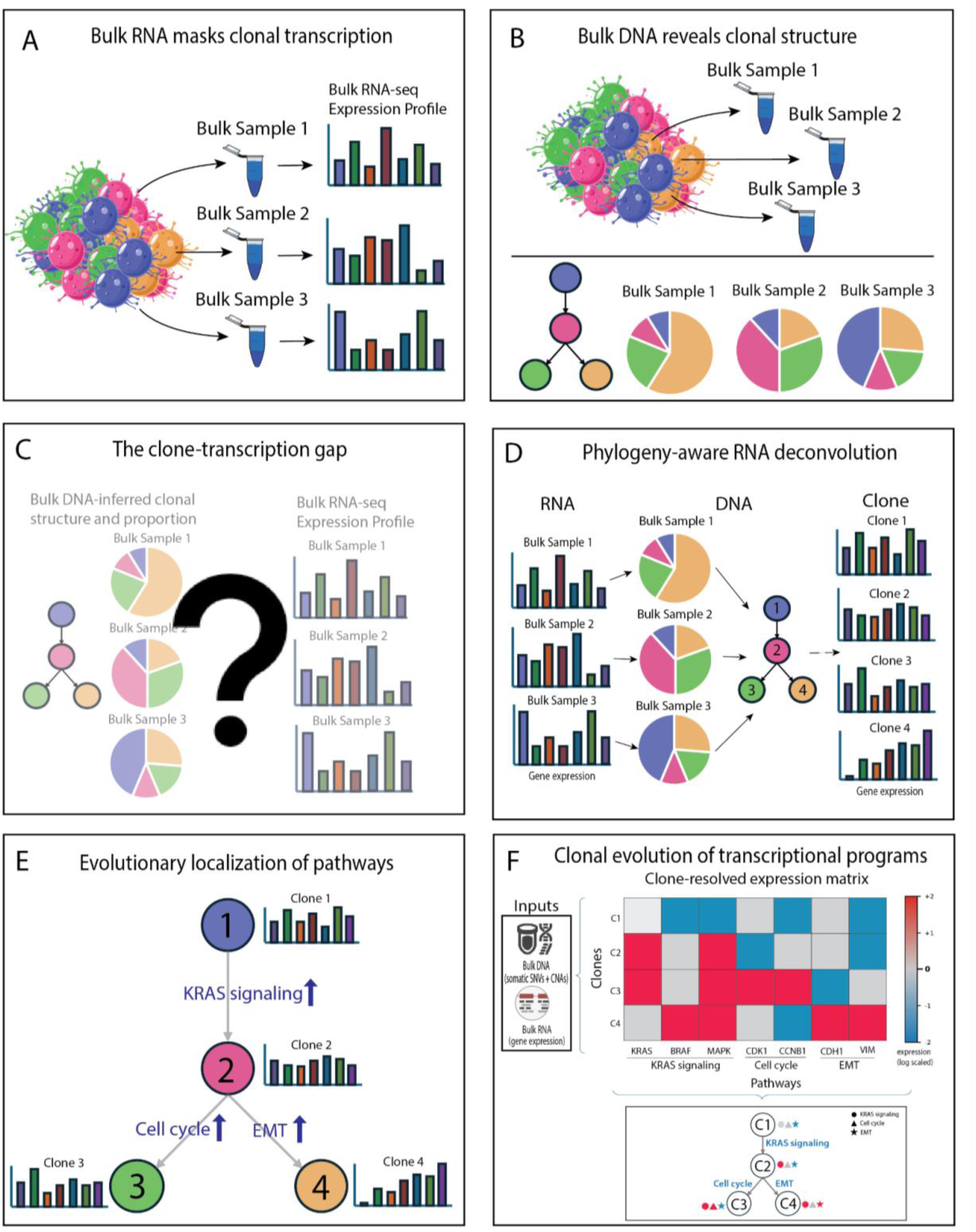
Overview of PICTographPlus for clone-resolved transcriptomic inference from bulk sequencing. (A) Bulk RNA-seq measures a composite transcriptional profile arising from multiple tumor clones, obscuring clone-specific gene expression programs. (B) In contrast, bulk DNA sequencing enables inference of clonal composition and evolutionary relationships across samples. (C) The resulting disconnect between DNA-inferred clonal structure and bulk RNA expression has limited reconstruction of clone-resolved transcriptional programs. (D) PICTographPlus bridges this gap by integrating DNA-inferred clonal phylogenies with bulk RNA-seq using a regularized mixture model that aligns DNA-derived clone proportions with observed expression to infer clone-specific transcriptomes. (E) By modeling transcriptional changes along phylogenetic edges, the method localizes pathway gains and losses to specific evolutionary transitions rather than to individual bulk samples. (F) PICTographPlus outputs clone-resolved gene expression and gene set activity mapped onto tumor evolutionary trees, enabling retrospective and cohort-scale analyses from widely available bulk sequencing data.

### The two genomic modalities meet in a single optimization

Given the DNA-derived clone proportion matrix Π (samples x clones) and the bulk gene expression matrix Y (samples x genes), PICTographPlus infers a clone-by-gene expression matrix X by minimizing a two-term objective:

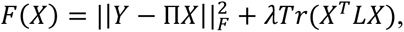

where the first term is a reconstruction loss that fits the clone-weighted mixture to each sample, and the second term penalizes large expression differences between clones connected in the regularization graph. L is the Laplacian matrix of this graph, and lambda is a regularization parameter controlling the strength of smoothing. In the general formulation, L can encode the clonal tree (Laplacian regularization, which assumes that phylogenetically adjacent clones have more similar expression) or a star graph in which all clones are connected only to the root (star regularization, which applies uniform shrinkage without imposing phylogenetic distance relationships). We evaluated seven regularization models (plain, adaptive, adaptive_v2, plain_debiased, fused_ew, elastic_net, and tree_delta) that differ in how this regularization is applied (Methods). Systematic topology sensitivity analysis (see below) showed that star regularization consistently matched or outperformed tree-structured. We therefore use star regularization for deconvolution and reserve the clonal phylogeny for its unique interpretive role: mapping recovered expression changes onto specific evolutionary branches.

To map transcriptional changes onto evolutionary transitions, we compute gene-level log2 fold changes along each edge (child clone vs parent clone) and rank genes by their edge-specific fold changes. We then perform gene set enrichment analysis (GSEA) on these ranked lists for curated gene set collections, thereby localizing gene set gains and losses to specific clonal branches (Figure 1D-E). The output of PICTographPlus includes (i) an inferred clonal tree, (ii) per-sample clone fractions, (iii) clone-specific expression profiles, and (iv) edge-localized gene set programs (Figure 1F).

### Benchmarking using single-cell RNA with matched single-cell DNA sequencing data

Validating clone-level inference requires a ground-truth dataset where DNA-defined clone membership and clone-specific transcriptomes are jointly observed in the same cells, a measurement combination that has historically been impossible at scale. The wellDR-seq protocol recently overcame this limitation by simultaneously profiling the genome and transcriptome of individual single cells^18^. We used this technology’s resolution to construct a controlled validation framework for PICTographPlus, working from patient P7 of an ER+ breast cancer cohort (Figure 2A). Because clone membership (from scDNA) and expression (from scRNA) are measured in the same cells, pseudo-bulk profiles aggregated within each clone represent ground-truth clone-level expression that PICTographPlus aims to recover from bulk mixtures. P7 scDNA identified one diploid normal clone (c1) and four tumor clones (c2, c4, c7, c11), plus a rare subclone c13 (n = 16 cells) carrying only the chr16q loss. Because c13 shares its only copy-number event (chr16q loss) with c11 and its exclusion produced an artifactual polyclonal topology with two independent branches from the diploid root, we merged c13 cells into c11, yielding the trunk clone c11_c13 (n = 176 cells) and a biologically correct monoclonal tree (Figure 2B). The P7 dataset also provides an opportunity to examine, with cellular resolution, how related clones of a single tumor differ transcriptionally, a question that has been hard to address with conventional single-modality methods.

**Figure 2.**
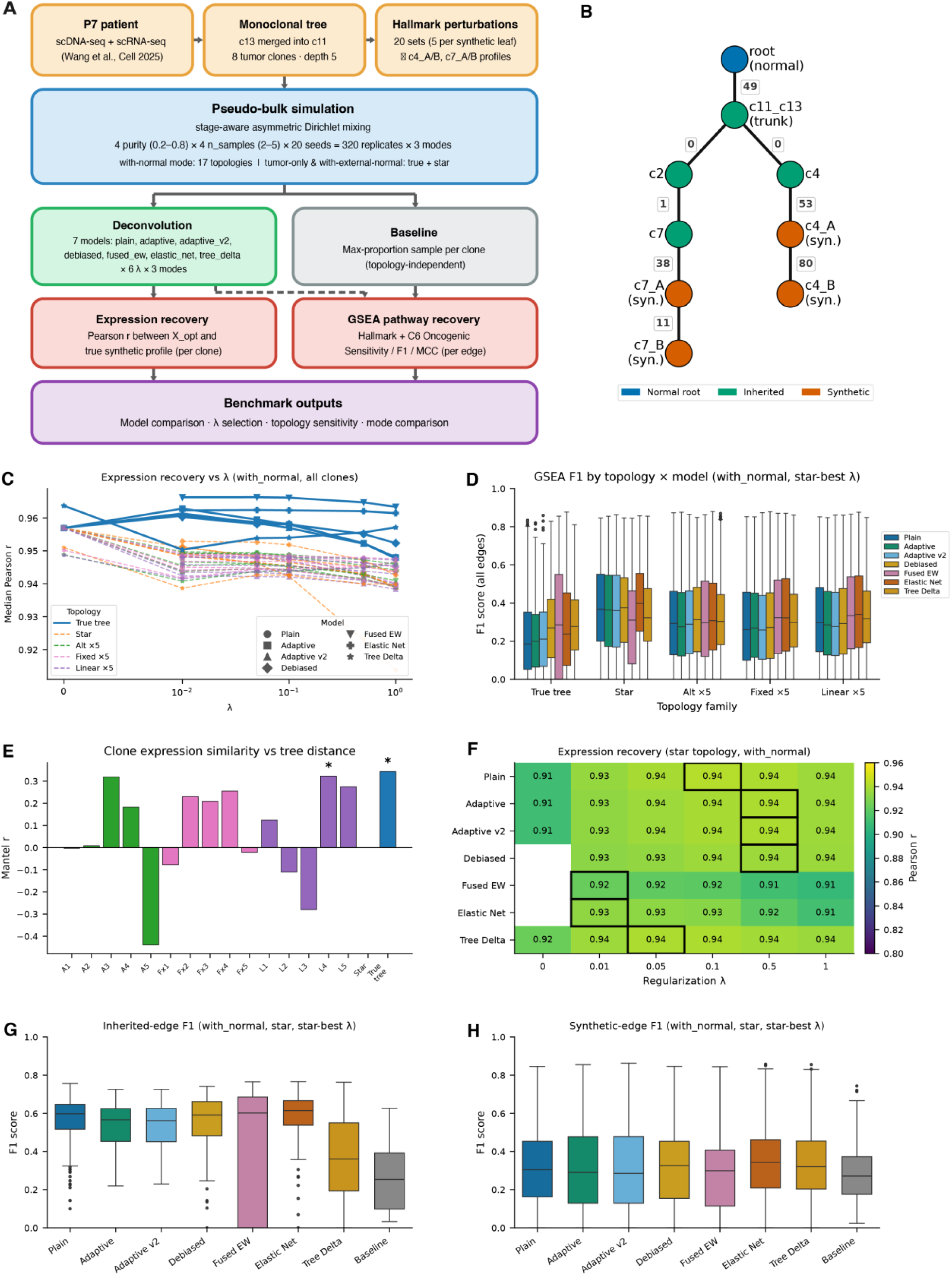
Benchmarking PICTographPlus on a corrected deep P7 clonal tree. (A) Benchmark design; (B) corrected P7 tree with 1 normal root, 4 inherited tumor clones, 4 synthetic clones, edge-labeled with ground-truth pathway counts. (C) Expression recovery vs λ across topology families (with-normal). (D) GSEA F1 across topology families at each model’s star-best λ. (E) Clone-expression similarity vs tree distance: Mantel r for each topology; only the true tree achieves one-sided p ≤ 0.05. (F) Heatmap of expression recovery on the star topology across λ (black outlines = best λ per model). (G) GSEA F1 on inherited edges (star). (H) GSEA F1 on synthetic edges (star).

To stress-test the method under a deeper clonal hierarchy and to enable controlled assessment of topology sensitivity, we extended this corrected tree by treating c7 and c4 as internal nodes and growing synthetic clones along two independent lineages, with expression profiles generated by multiplicatively perturbing the parent clone’s pseudo-bulk profile using fold changes derived from the Hallmark gene sets (MSigDB). Five gene sets were assigned uniquely to each synthetic clone, with no overlap across clones. The resulting tree comprises K = 8 tumor clones with a maximum depth of 5 edges, providing a richer topology structure for systematic evaluation (Figure 2B; Methods). We generated 320 replicate pseudo-bulk datasets by mixing clone-level expression profiles using a stage-aware asymmetric Dirichlet sampling scheme spanning four tumor purity levels (0.20, 0.40, 0.60, 0.80), four numbers of tumor bulk samples (2, 3, 4, 5), and 20 random seeds per condition (Methods).

### The correct clonal phylogeny does not improve deconvolution over topology-agnostic regularization

We evaluated whether providing the true (DNA-derived) clonal phylogeny to PICTographPlus improved deconvolution accuracy over alternative topologies. In with-normal mode we evaluated all seven models under 17 tree topologies: the true tree, a star topology (all clones equidistant from the root), five randomly permuted alternatives (alt x 5), five fixed-structure permutations (fixed x 5), and five linear chains (linear x 5) (Methods).

### Expression recovery

The true clonal phylogeny provided a consistent but modest advantage in per-clone expression recovery over all tested alternative topologies across all models and regularization strengths (Figure 2C, Figure S1). ΔPearson 𝑟 (true tree minus alternative-family mean) was positive throughout, but magnitudes were small: at the star-best 𝜆, advantages ranged from +0.007 to +0.027 for synthetic clones and +0.009 to +0.044 for inherited clones.

Importantly, when topology advantage is quantified as the per-replicate difference between the true tree and the mean across alternative topology families (star, alt, fixed, linear), all seven regularization models showed a small but statistically significant positive expression-recovery advantage at λ = 0.1 (synthetic clones, 320 paired replicates, with-normal mode). Hodges-Lehmann estimates of the mean per-replicate ΔPearson 𝑟 (true tree − mean of alternative topology families) ranged from +0.012 (plain_debiased; 95% CI [+0.011, +0.013]) to +0.024 (fused_ew; 95% CI [+0.021, +0.026]) across the seven models, with all paired Wilcoxon signed-rank tests significant at p < 0.001 (Bonferroni-corrected; per-model values in Table S1). The star topology yielded the smallest gap of the four alternative families in every model, confirming it as the closest uninformative approximation to the true tree.

### GSEA pathway recovery

Across all evaluable edges, the star topology outperformed the true tree for all seven models at their star-best 𝜆 in with-normal mode (Figure 2D). Star median F1 ranged from 0.31 to 0.40 versus 0.19 to 0.29 for the true tree, with the largest gaps in the Laplacian-family models (plain = −0.18, adaptive =−0.16, adaptive_v2 =−0.15) and elastic_net (−0.16). The Laplacian collapse was driven primarily by the inherited root edge: the deep true tree heavily shrinks the trunk log-fold change via accumulated chain coupling, whereas the star’s uniform depth of 1 avoids this penalty (true-tree median F1 on inherited edge: 0.07-0.09 vs. 0.56-0.60 under the star) (Figure S2). The star also outperformed the true tree on synthetic edges alone (gap 0.05-0.14), indicating an independent benefit of shallower topology. These results motivated adopting the star as the topology for all manuscript figures.

#### Inter-clone expression similarity does not scale with phylogenetic distance

The benchmark also produced a biological observation about inter-clone divergence in this tumor. We computed pairwise expression similarity (Pearson r) between all 8 tumor clone profiles and tested whether similarity decreases with tree distance under each of the 17 topology variants. Inter-clone expression similarities were compressed into a narrow range (0.77-0.99 Pearson r) that did not track phylogenetic distance: clones distant on the true tree could be nearly as similar as adjacent clones (Figure S3, Table S2).

Spearman rank correlation between expression similarity and true tree distance was rho = +0.34, the highest value across all 17 topologies but only marginally significant (Figure 2E). A Mantel permutation test (9,999 permutations, one-sided) confirmed a positive but weak association for the true tree (Mantel 𝑟 = 0.34, 𝑝 = 0.04, exact permutation test); the true tree was the only topology achieving Mantel significance, but several randomly scrambled alternatives came close (e.g., alt_3: 𝑟ℎ𝑜 = 0.32; linear_4: 𝑟ℎ𝑜 = 0.32; Figure 2E, Table S3). The weak correlation between phylogenetic distance and expression similarity reflects a fundamental property of cancer biology: inter-clone transcriptomic differences are sparse and pathway-specific, embedded in a large, shared background (> 95% common transcriptome across clones of the same tumor). Unlike species-level phylogenetics where mutations accumulate proportionally over time genome-wide, a single clonal transition in a tumor typically affects only a small number of pathways, while the remaining transcriptome is invariant. As a result, a clone at tree distance 1 (differing by one mutation-associated program) can be as globally similar to a distant clone as to its direct parent. Mechanistically, this reflects a property of tumor evolution that has no analog in species phylogenetics: a single subclonal driver event can reprogram a small set of pathways while leaving the bulk transcriptome unchanged, so phylogenetic distance and transcriptomic divergence are decoupled.

### Chain smoothing on deep trees dilutes localized edge signals

We identified a second mechanism by which the true phylogeny degrades deconvolution: chain smoothing in Laplacian-regularized models. The Laplacian penalty couples intermediate nodes through a joint inverse, attracting each clone’s expression toward both parent and children. On a deep tree, a perturbation on a specific edge (e.g., c7 → c7_A) is smeared across the entire chain root → c11_c13 → c2 → c7 → c7_A → c7_B. The fold-change contrast that GSEA requires on the c7 → c7_A edge is diluted, reducing sensitivity on that edge (Figure S2).

On the star topology, by contrast, every clone is directly connected only to the root: there are no chains and no cross-node propagation. A perturbation on c7_A remains localized to the c7_A - root edge contrast. This is why the star topology outperforms the true tree for Laplacian models on synthetic edges: not because it encodes better biological structure, but because it avoids the chain-propagation artifact that the true tree introduces. The tree_delta model, which parameterizes clone expression as cumulative edge deltas and penalizes each delta independently (rather than as a global Laplacian), showed the smallest topology-dependent gap in both expression recovery and pathway recovery on synthetic edges, consistent with its edge-separable design (Figure S2).

### The phylogeny’s role is interpretive, not regularizing

These results establish a key principle: the clonal phylogeny’s value lies in defining edges along which transcriptomic transitions are interpreted, not in improving deconvolution through tree-structured regularization. The Mantel analysis shows the true tree’s distance-similarity signal is too weak to serve as a useful regularization prior, and topology benchmarking confirms the star topology performs as well as or better than the true tree for most models. The true tree remains essential for interpretation: it specifies which clone pairs define each edge, identifies directionality of change, and allows pathway enrichment to be assigned to specific evolutionary transitions. The regularization prior is separable from this interpretive role, and star regularization is the empirically better choice. We therefore use star regularization for all subsequent analyses.

### PICTographPlus achieves high expression recovery across all regularization models

In with-normal mode using the star phylogeny, all seven regularization models recovered clone expression profiles with mean Pearson 𝑟 ranging from 0.92 (fused_ew; 95% CI [0.90, 0.94]) to 0.94 (plain; 95% CI [0.93, 0.95]) across the seven models at their optimal regularization strength (Table S4, Figure 2F). At 𝜆 = 0 (no regularization), recovery was noticeably lower for plain, adaptive, and adaptive_v2, confirming that regularization provides meaningful benefit for expression recovery even when the magnitude of the optimal lambda is small.

Inherited (real single-cell) clones and synthetic clones showed distinct recovery profiles that varied substantially across deconvolution modes (Figure S4). Synthetic clones achieved consistently high mean Pearson 𝑟 (ranging from 0.94 [fused_ew; 95% CI 0.92, 0.95] to 0.95 [plain; 95% CI 0.94, 0.96] across models), reflecting the strong, localized fold-change signal embedded in the synthetic perturbations. Inherited clone recovery was markedly mode-dependent: including a matched normal reference (with-normal mode) yielded the highest and most uniform recovery (Pearson 𝑟 ranging from 0.91 (fused_ew; 95% CI [0.89, 0.93]) to 0.93 (adaptive/tree_delta; 95% CI [0.92, 0.95]) across models), whereas tumor-only mode showed the largest model spread (0.73-0.93) and lowest overall values. With-external-normal performance was intermediate (0.83-0.93), with fused_ew and elastic_net still reduced relative to the matched-normal condition. In tumor-only mode, Laplacian-family models (plain, adaptive, adaptive_v2) recovered only Pearson 𝑟 = 0.57 for inherited clones at 𝜆 = 0 and required regularization to reach ≥ 0.93, highlighting the critical role of both a normal reference and adequate regularization for faithful reconstruction of inherited clone expression.

### The normal reference is critical for root-adjacent edges but not for deeper clone transitions

We assessed how including a normal reference sample affects pathway recovery across tree edges of different depths (Figures S5-6). The impact is highly depth dependent. For the root-adjacent edge (edge_0_1: root → c11_c13, depth 1, 49 ground-truth significant pathways), the normal reference provided a substantial and consistent benefit across deconvolution models. At the star-best 𝜆, median F1 on this edge ranged from 0.56-0.62 in with-normal mode versus 0.22-0.28 in tumor-only mode for six of the seven models (approximately 2.0-2.6-fold improvement; Figure S5). Tree_delta was the exception, achieving a lower median F1 of 0.36 in with-normal mode due to the tree-structured penalty shrinking the trunk log fold-change under the uninformative star prior. The benefit of the normal reference on this edge is mechanistically direct: the normal sample pins the root clone expression to a known reference, anchoring the log fold-change computed on this edge. Without this anchor, the absolute expression scale of the root clone is poorly constrained, making the root-to-trunk LFC ambiguous.

For the four synthetic edges at depths 3-5, the normal reference provided minimal additional benefit at the star-best 𝜆. Pooled across all seven models, median F1 values were comparable between tumor-only and with-normal modes: edge_3_4 (depth 3) 0.30 vs 0.32; edge_4_5 (depth 4) 0.32 vs 0.36; edge_6_7 (depth 4) 0.31 vs 0.33; edge_7_8 (depth 5) 0.24 vs 0.24 (Figure S6). This pattern reflects the structure of the deconvolution problem: on edges between two tumor clones, the GSEA signal is the LFC between child and parent expression, a relative quantity that does not directly depend on the absolute root-scale anchor. The normal sample constrains the tumor-root pair but not the relative differences among tumor clones. Only on the root-to-trunk edge does anchoring the root directly stabilize the relevant LFC. All further analyses are conducted in with-normal mode to ensure optimal recovery of the root-adjacent edge and overall expression fidelity.

### An independent external normal reference provides intermediate performance

To assess whether the choice of normal reference affects deconvolution accuracy, we compared with-normal mode (c1, the within-dataset diploid normal, n = 297 cells) to with-external-normal mode (population-average normal from a breast cell atlas, 456,493 cells)^26^ (Methods).

Expression recovery and GSEA pathway F1 in with-external-normal mode were consistently intermediate between tumor-only and with-normal across all models (Figures S4-6). This partial recovery suggests that even an imperfectly matched external normal provides useful deconvolution grounding, while confirming that the matched within-dataset normal provides the most informative anchor.

### Deconvolution outperforms baseline

On the most informative inherited edge (edge_0_1: root → c11_c13), deconvolution models substantially outperformed the max-proportion baseline in GSEA pathway recovery (Figure 2G, Figure S5). At their star-tree-best 𝜆, six of the seven models (plain, adaptive, adaptive_v2, plain_debiased, fused_ew, and elastic_net) achieved median F1 of 0.56-0.62 and median sensitivity of 0.63-0.69, compared to a baseline median F1 of 0.25 and sensitivity of 0.18 (Figure 2G). Elastic_net (𝜆 = 0.01) achieved the highest median F1 on this edge (0.62; sensitivity 0.69), followed closely by plain (0.60; 0.68) and fused_ew (0.60; 0.63). The baseline’s poor performance on this edge reflects a fundamental limitation of the no-deconvolution approach: without explicitly separating each clone’s contribution, the expression profile of the maximum-proportion sample is dominated by the most abundant clone and confounded by all others present. Deconvolution recovers the specific transcriptional program associated with the root-to-trunk transition, which the baseline cannot access directly. Three edges (c11_c13 → c2, c11_c13 → c4 and c2 → c7) had zero or one ground-truth significant pathways and were therefore excluded from sensitivity and F1 computation.

On the four synthetic edges (depths 3-5), deconvolution models achieved meaningful but more modest pathway recovery relative to the inherited root edge (Figure 2H). Pooled across all four synthetic edges, the best-performing regularization models at their star-tree-best λ achieved median F1 of 0.29-0.34 (Table S5), versus a baseline median F1 of 0.27. Performance varied markedly across the four synthetic edges and was shaped by both depth and ground-truth pathway count: edge_4_5 (depth 4, GT = 80 pathways) achieved the highest pooled median F1 (∼0.36), followed by edge_6_7 (depth 4, GT = 38, ∼0.33) and edge_3_4 (depth 3, GT = 53, ∼0.30). The deepest edge, edge_7_8 (depth 5, GT = 11), showed the lowest recovery (∼0.23), reflecting both the progressive dilution of deep subclonal signals in the bulk mixture and the reduced number of evaluable ground-truth pathways at greater tree depth.

### Linking clonal evolution and transcriptional programs in NSCLC

We applied PICTographPlus to a TRACERx non-small cell lung cancer case (CRUK0004) to ask what clone-resolved transcriptional analysis can reveal about the trajectory of a primary tumor through surgery, systemic therapy, and metastatic relapse, questions that bulk RNA region-level analyses leave unresolved. CRUK0004 is a Stage IIB NSCLC treated surgically and later relapsing after vinorelbine^19–21^, with bulk DNA and RNA available from four primary tumor regions and a lymph node metastasis collected at progression, with matched normal adjacent lung tissue. PICTographPlus inferred a six-clone evolutionary tree in which a TP53 p.Glu68Ter nonsense (stop-gain) mutation marked the founding malignant clone and subsequent branching events introduced additional drivers, including an EGFR-mutant branch, a SMAD4/NOTCH1-mutant branch, and a terminal EGFR-amplified metastatic clone (Figure 3A-C, Figure S7A).

**Figure 3.**
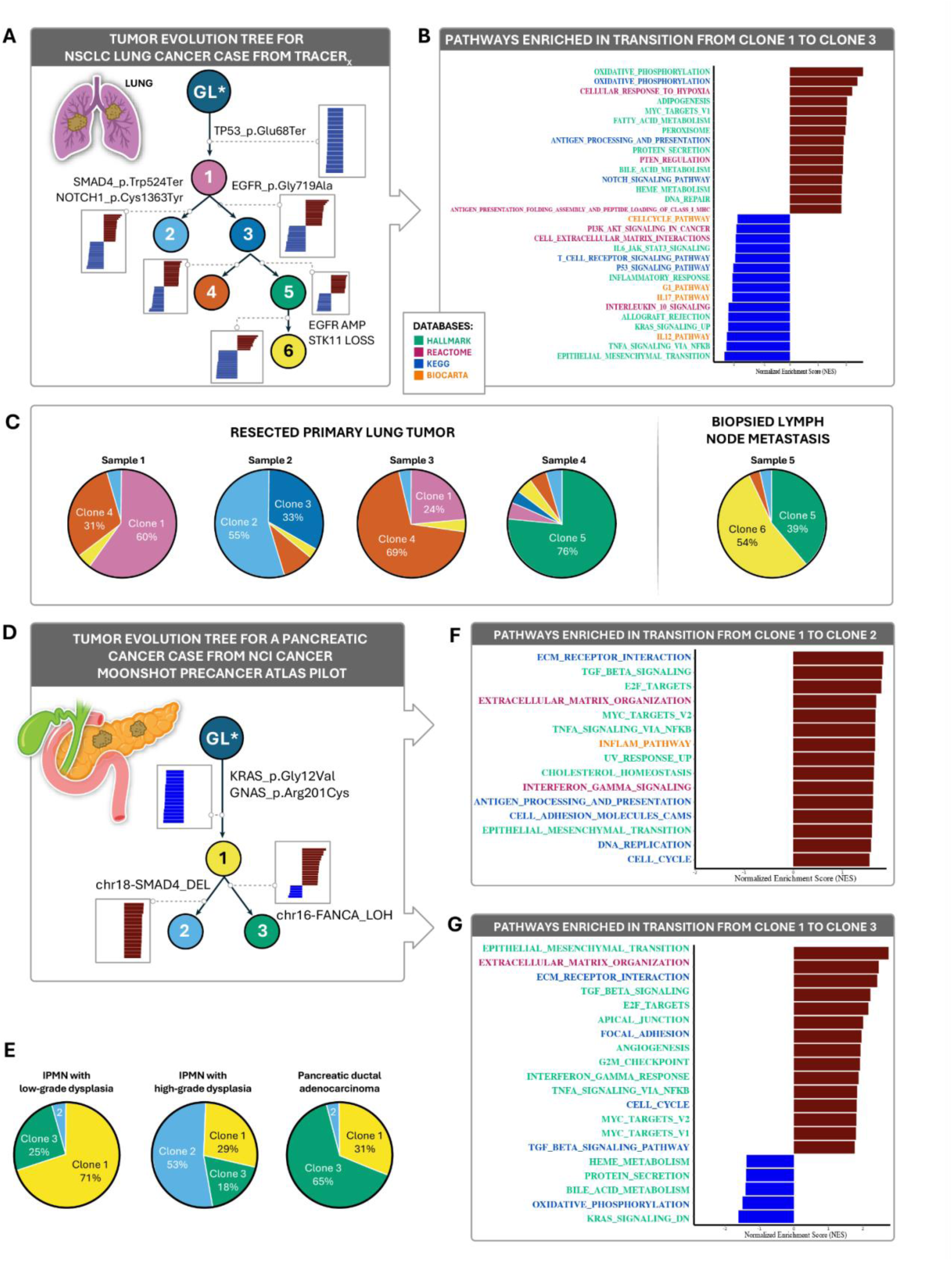
Clone-resolved evolution and gene set shifts in lung and pancreatic tumors inferred by PICTographPlus. (A) CRUK0004 clonal tree; truncal TP53 p.Glu68Ter (nonsense/stop-gain), SMAD4/NOTCH1 mutation on Clone 2, EGFR mutation on Clone 3, terminal EGFR amplification + STK11 loss on metastatic Clone 6. (B) NES bar plot for Clone 1 to Clone 3. (C) Subclone composition across 5 tumor regions. (D) MK74 three-node tree with KRAS G12V/GNAS R201C trunk, SMAD4-deleted Clone 2, FANCA-LOH Clone 3. (E) Stage-specific subclone proportions across LGD IPMN, HGD IPMN, invasive PDAC. (F,G) Edge-wise NES for the SMAD4 and FANCA branches respectively.

Edge-level GSEA revealed distinct transcriptional programs at each transition. The germline-to-Clone 1 edge showed broad suppression relative to normal lung, spanning oxidative phosphorylation, MHC class I antigen presentation, MYC targets, TGF-β signaling, and multiple immune and metabolic programs (Figure S7B). The Clone 1-to-Clone 2 transition showed coordinated metabolic activation (oxidative phosphorylation, glycolysis, fatty acid metabolism, hypoxia response, MYC targets, DNA repair) alongside concurrent immune suppression (TNFα/NFκB, EMT, inflammatory response, IL6-JAK-STAT3, T-cell receptor signaling) (Figure S7C). The Clone 1-to-Clone 3 edge showed a closely parallel pattern (Figure 3B), indicating a shared metabolic-activation and immune-downregulation program on both intermediate branches. Further downstream, Clone 3 to Clone 4 acquired a strong immune/proliferation program (allograft rejection, PD-1 signaling, E2F targets, G2/M checkpoints, MYC targets, PI3K-AKT-mTOR) (Figure S7D), while the sister Clone 3-to-Clone 5 edge combined interferon/EMT/proliferative programs with suppression of senescence, mTOR, and p53 pathway activity (Figure S7E). Finally, the terminal Clone 5-to-Clone 6 edge, corresponding to the EGFR-amplified metastatic clone, was dominated by widespread loss of metabolic and stress-response programs (mTORC1, oxidative phosphorylation, glycolysis, hypoxia response, DNA replication) together with modest upregulation of T-cell/adaptive-immune modules (PD-1 signaling, IL12, IL7, T-cell receptor signaling) (Figure S7F). In this patient, PICTographPlus localizes a coordinated metabolic-activation/immune-suppression program (oxidative phosphorylation, glycolysis, fatty-acid metabolism, MYC targets, DNA repair, with concurrent suppression of TNFα/NFκB, EMT, IL6-JAK-STAT3, and T-cell receptor signaling) to a specific intermediate clonal transition, and a distinct T-cell/PD-1-associated transcriptional state to the terminal EGFR-amplified metastatic clone that emerged after therapy. The clinical implication is that the immune-cold state characterizing the post-therapy metastasis is not a property of the metastasis-as-a-whole but of a specific clonal lineage, a distinction that bulk-region analyses cannot resolve and that has implications for targeting therapy-resistant lineages. Whether this pattern recurs across NSCLC will require systematic application to larger TRACERx-scale cohorts.

### Clonal evolution from pancreatic precursor to invasive PDAC

Whether the transcriptional programs that drive precursor-to-invasive transition in pancreatic cancer share a single trajectory or diverge across lineages has been a difficult question to address, invasive carcinomas and their precursor lesions are typically compared as bulk pairs, averaging over any clonal substructure within each lesion. We applied PICTographPlus to pancreatic case MK74 from the NCI Cancer Moonshot Precancer Atlas Pilot^22^, which provides laser-capture-microdissected (LCM) bulk RNA and DNA from a low-grade IPMN, a high-grade IPMN, and invasive PDAC from the same patient. PICTographPlus reconstructed a three-clone tree in which a common ancestor (Clone 1) bearing KRAS G12V and GNAS R201C predominated in the LGD IPMN and seeded two descendant lineages (Figure 3D-E). The SMAD4-deleted descendant (Clone 2) expanded in the HGD IPMN and was associated with a coordinated proliferative and stromal-remodeling program: upregulation of ECM-receptor interaction, TGF-β signaling, E2F targets, MYC targets, DNA replication, cell cycle, and EMT (Figure 3F). The FANCA LOH descendant (Clone 3), dominant in invasive PDAC, showed an even more pronounced EMT/ECM-remodeling and cell-cycle programs, with concurrent suppression of oxidative phosphorylation and bile acid/heme metabolism (Figure 3G). The implication is that within a single patient, two clonal lineages can pursue parallel but transcriptionally distinct routes to malignancy: one expanding proliferatively in high-grade precursors after SMAD4 loss, another acquiring homologous recombination (HR) deficiency and EMT/ECM remodeling on the path to invasion. Region-level comparisons cannot resolve this because each lesion is itself a clone mixture; only clone-resolved analysis separates the two trajectories. This biological observation generalizes a possibility long suspected in IPMN-PDAC progression studies19 but previously inaccessible from bulk data.

### Organ-specific adaptation in a rapid-autopsy PDAC with multi-organ metastases

The most distinctive feature of metastatic cancer is organ specificity: tumor cells that have left the primary site adapt transcriptionally to the niche they colonize. Whether sister metastatic clones diverging from a shared trunk pursue parallel adaptive strategies or converge on a common metastatic program is a central question in cancer biology. We applied PICTographPlus to PDAC case PCSI0380 from the PanCuRx rapid-autopsy series^23^, comprising a primary PDAC, a lymph-node metastasis, and a liver metastasis from a single patient, with matched normal DNA from skeletal muscle. PICTographPlus inferred an architecture in which a truncal KRAS G12D/TP53 R175H/ARID1A-mutant Clone 1 gives rise to Clone 2 (dominant in the liver metastasis, 95%) and Clone 3 (present in both primary, 43%, and lymph node metastasis, 25%) (Figure 4A–B). Edge-level GSEA comparing Clone 2 to Clone 1 revealed a strong shift toward inflammatory and liver-adapted programs: TNFα/NFκB signaling, hypoxia, coagulation, complement, IL6-JAK-STAT3, interferon-γ response, xenobiotic and bile acid metabolism, and elevated KRAS-up output, with relative attenuation of E2F targets, MYC targets, DNA replication, cell cycle, and oxidative phosphorylation (Figure 4C). The liver-metabolic signal (xenobiotic/bile acid/heme) likely partly reflects hepatocyte transcripts in the bulk mixture, but the inflammatory/KRAS-up/cytokine programs are less plausibly explained by contamination.

**Figure 4.**
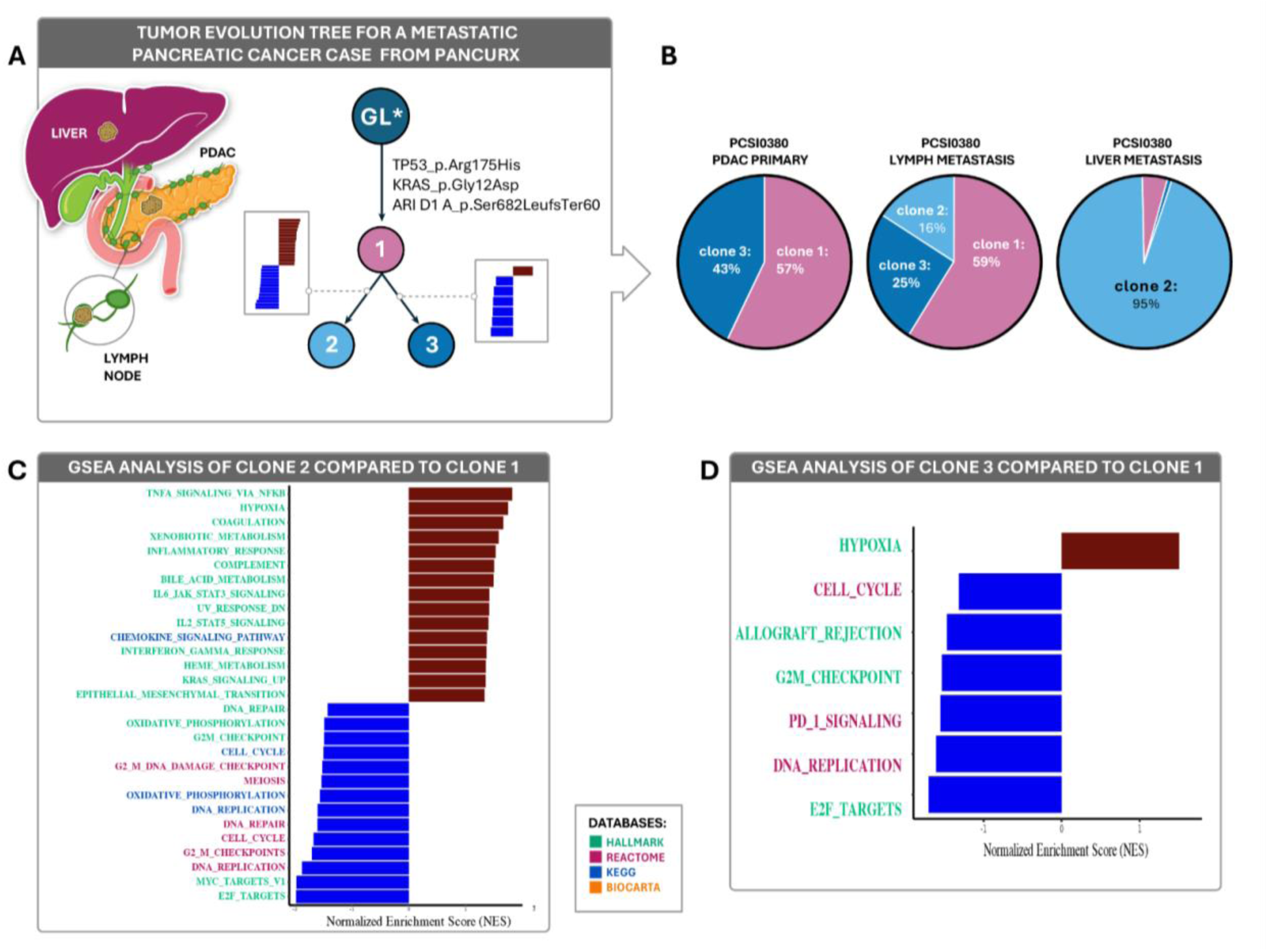
Distinct microenvironmental adaptations of two daughter clones in PanCuRx case PCSI0380. (A) Clonal evolutionary tree rooted at TP53 R175H/KRAS G12D/ARID1A-mutant Clone 1. (B) Clone proportions across primary, lymph node, and liver metastasis. (C) GSEA of Clone 2 vs Clone 1 (liver-dominant): upregulation of inflammatory/stromal programs; downregulation of replication, cell cycle, OXPHOS. (D) GSEA of Clone 3 vs Clone 1 (primary/LN): single enrichment (hypoxia) with coordinated suppression of cell cycle, G2/M, allograft rejection, PD-1, replication, E2F.

GSEA of Clone 3 versus Clone 1 revealed only a single significant enrichment (hypoxia), with coordinated depletion of E2F targets, DNA replication, G2/M checkpoint, cell cycle, and immune programs including PD-1 signaling and allograft rejection (Figure 4D). In this patient, PICTographPlus resolves the question posed above in a specific direction: sister metastatic clones diverge from a shared KRAS/TP53/ARID1A trunk into two transcriptionally distinct, organ-tuned states rather than converging on a single metastatic program. The liver-dominant clone adopts an inflamed, liver-metabolic, genomically permissive state; the primary-and-lymph-node-dominant clone adopts a hypoxic, low-proliferation, immune-quiet state. Site-level bulk RNA comparisons would average these together and miss both states. Whether this pattern of clone-by-organ divergence recurs across rapid-autopsy PDAC cohorts is now an answerable question; the framework brings such cohorts within reach of clone-resolved analysis.

## Discussion

PICTographPlus addresses a gap between two well-established genomic data modalities: bulk DNA-seq, which has become the dominant tool for inferring tumor evolutionary architecture, and bulk RNA-seq, which has become the dominant tool for profiling tumor transcriptional state, but which until now has not been bridgeable to the clone-resolved transcriptional programs that DNA-defined clones must possess. By framing this bridge as a joint inference problem, DNA-derived clone proportions and phylogeny supplying the structural constraints, bulk RNA supplying the observed mixture, and a regularized least-squares decomposition supplying the latent clone-specific expression matrix, the framework makes clone-resolved transcriptional analysis tractable for the cohorts where it has historically been out of reach: multi-region, longitudinal datasets that are now numerous but for which matched single-cell profiling is not available.

Across three cases, PICTographPlus illustrated its ability to resolve lineage-specific programs that are invisible to bulk comparisons. In the NCI Precancer Atlas IPMN case, two sister clones diverged from a KRAS/GNAS trunk: the SMAD4-deletion clone (clone 2) expanded substantially in abundance between low- and high-grade dysplasia, with its transition enriched for proliferative and ECM programs, whereas the FANCA-LOH clone (clone 3) acquired a dominant epithelial–mesenchymal-transition program with coordinated suppression of oxidative-metabolism gene sets. In the rapid-autopsy PDAC, sister clones from a shared malignant trunk diverged into an inflamed, liver-adapted state versus a hypoxic, immune-quiet state. In TRACERx, intermediate branches carried coordinated metabolic and immune programs, while the terminal EGFR-amplified clone shifted toward a T-cell/PD-1 signature after therapy. These clone-resolved features, opposing directions on sister edges, driver-program co-localization, and divergent sister states, are presented here as the kinds of patterns deconvolution can recover from individual patients; their generalizability across larger cohorts is not a claim of this work and remains an open question.

A key methodological design choice in PICTographPlus is that the clonal phylogeny enters in two distinct roles: as DNA-derived constraints on clone proportions and tree structure, and as an interpretive scaffold for projecting recovered transcriptional differences onto evolutionary branches. It does not enter as a regularization prior on the deconvolution itself — a choice we initially expected to be a limitation, but that our systematic benchmark established as the empirically better one. The reason is biological as much as numerical: inter-clone transcriptional divergence is sparse and pathway-specific, so penalties that smooth expression along tree edges (as a phylogeny-aware prior would) blur the very differences the method is meant to recover.

Consistent with this, although a Mantel test identified the true phylogeny as the only topology whose pairwise distances track scRNA-derived clone-expression similarity, that correlation is itself weak and only marginally significant (Spearman ρ = 0.34, with permuted-topology alternatives close behind), and a structureless star topology recovered pathway-level differences as well as or better than the true tree across all regularization models, so we adopt the star as the evaluation topology throughout. Beyond topology, the benchmark identified the two choices that matter most in practice: including a matched normal reference, which improved pathway recovery on root-adjacent edges roughly 2–2.6 fold over tumor-only analysis (with diminishing benefit at deeper tumor-to-tumor edges), and adequate regularization. Full quantitative results, including overall expression-recovery accuracy (mean Pearson r ≥ 0.92), per-model rankings, and the mode-dependence of inherited-clone recovery, are reported in Results.

PICTographPlus has limitations. It depends on upstream clonal reconstruction and clone-fraction estimates, which remain challenging and model-dependent; mis-specified trees may mis-assign transcriptional changes to specific branches. Bulk RNA-seq conflates tumor and non-tumor compartments, so gene-set shifts involving immune or stromal signatures likely reflect coordinated changes in both malignant and microenvironmental cells, and our current implementation does not explicitly model these compartments or propagate uncertainty from DNA-based clonal inference to RNA-level estimates. Our quantitative benchmark is itself derived from the clonal structure of a single co-profiled patient, extended with synthetic clones; establishing these accuracy figures across diverse patients and tumor types will require broader ground-truth validation. More broadly, our applications focus on a small number of illustrative cases; systematic application to larger cohorts will be needed to identify recurrent clone-restricted programs, relate them to treatment and outcome, and evaluate whether branch-specific states have predictive or prognostic value.

Despite these caveats, PICTographPlus provides a general and scalable way to connect tumor evolution with transcriptional reprogramming using standard bulk DNA and RNA assays. By anchoring expression changes to defined edges of clonal trees, it enables “virtual” clone-level analyses in settings where single-cell profiling is impractical, including multi-region and rapid-autopsy studies of metastatic disease. In principle the same formulation extends beyond bulk RNA: because the model requires only a mixture with known clone proportions, scDNA-seq- or CNV-defined clones can be aggregated into clone-level pseudo-bulks, and spatial transcriptomic spots can be treated as samples with clone fractions inferred from co-registered DNA, a route to clone-resolved analysis in single-cell and spatial data that follows directly from the model’s assumptions, though we do not test it here. As metastatic cohorts expand, methods such as PICTographPlus should help establish clone- and transition-specific gene-set programs as a routine component of metastatic interpretation, enabling systematic discovery of organ-adapted programs and more precise links between evolutionary transitions, microenvironmental context, and clinical progression.

## Resource Availability

### Lead contact

Further information and requests for resources should be directed to and will be fulfilled by the lead contact, Rachel Karchin.

## Materials availability

This study did not generate new unique reagents.

## Data and code availability

- All data analyzed in this study are publicly available. The scDNA/scRNA co-profiled single-cell dataset used for benchmarking was generated by wellDR-seq and is available at GEO under accession GSE261713^18^. IPMN WES and RNA-seq data are available through dbGaP (phs002225.v3.p1). TRACERx NSCLC WES and RNA-seq data are available through EGA (EGAD00001009825 and EGAD00001009862). PanCuRx WGS and RNA-seq data are available through EGA (EGAD00001004551 and EGAD00001004548). Processed data generated in this study, including clone-level expression matrices, inferred clonal trees, and edge-level GSEA results, are publicly available at Mendeley Data (https://data.mendeley.com/datasets/cv66sgfcfn/2) as of the date of publication.
- All original code is publicly available at https://github.com/KarchinLab/pictographPlus and archived at Zenodo (DOI: https://doi.org/10.5281/zenodo.19896732). The code is released under the MIT license.
- Any additional information required to reanalyze the data reported in this paper is available from the lead contact upon request.

## Acknowledgements

We acknowledge funding from the NIH/NCI (U01CA271273 to R.K., L.D.W., E.J.F., L.T.K., J.L., K.N., P.B.N. and U54CA274371 to L.D.W., E.J.F.) and Break Through Cancer, Cambridge, MA (to R.K., L.D.W., E.J.F., L.T.K., J.L., A.B., P.B.N.).

This study makes use of data generated by The TRAcking Non-small Cell Lung Cancer Evolution Through Therapy (Rx) (TRACERx) Consortium and provided by the UCL Cancer Institute and The Francis Crick Institute. The TRACERx study is sponsored by University College London, funded by Cancer Research UK and coordinated through the Cancer Research UK and UCL Cancer Trials Centre. It also uses data generated by the PanCuRx Translational Research Initiative, a Canadian multicentre pancreatic cancer research program led by the Ontario Institute for Cancer Research (OICR) and collaborating institutions. The PanCuRx Translational Research Initiative is supported by the Ontario Institute for Cancer Research through funding provided by the Government of Ontario and by the Wallace McCain Centre for Pancreatic Cancer through the Princess Margaret Cancer Foundation, with additional support from the Terry Fox Research Institute, the Canadian Cancer Society Research Institute and the Pancreatic Cancer Canada Foundation.

We thank Jennifer E. Fairman, CMI, FAMI, for the illustration of Figures 3 and 4.

## Author contributions

J.L.: Conceptualization, Data Curation, Formal Analysis, Methodology, Software, Validation, Visualization, Writing – Original Draft, Writing – Review & Editing. Y.Y.: Software. K.N.: Writing – Review & Editing. Y.L.: Writing – Review & Editing. A.B.: Data Curation, Methodology. P.B.N.: Conceptualization, Data Curation, Investigation, Methodology, Validation. L.T.K.: Formal Analysis, Funding Acquisition, Writing – Review & Editing. E.J.F.: Conceptualization, Funding Acquisition, Investigation, Methodology, Supervision, Validation, Visualization, Writing – Original Draft, Writing – Review & Editing. L.D.W.: Investigation, Writing – Review & Editing. R.K.: Conceptualization, Formal Analysis, Funding Acquisition, Investigation, Methodology, Resources, Supervision, Writing – Original Draft, Writing – Review & Editing.

## STAR★Methods

### Key resources table

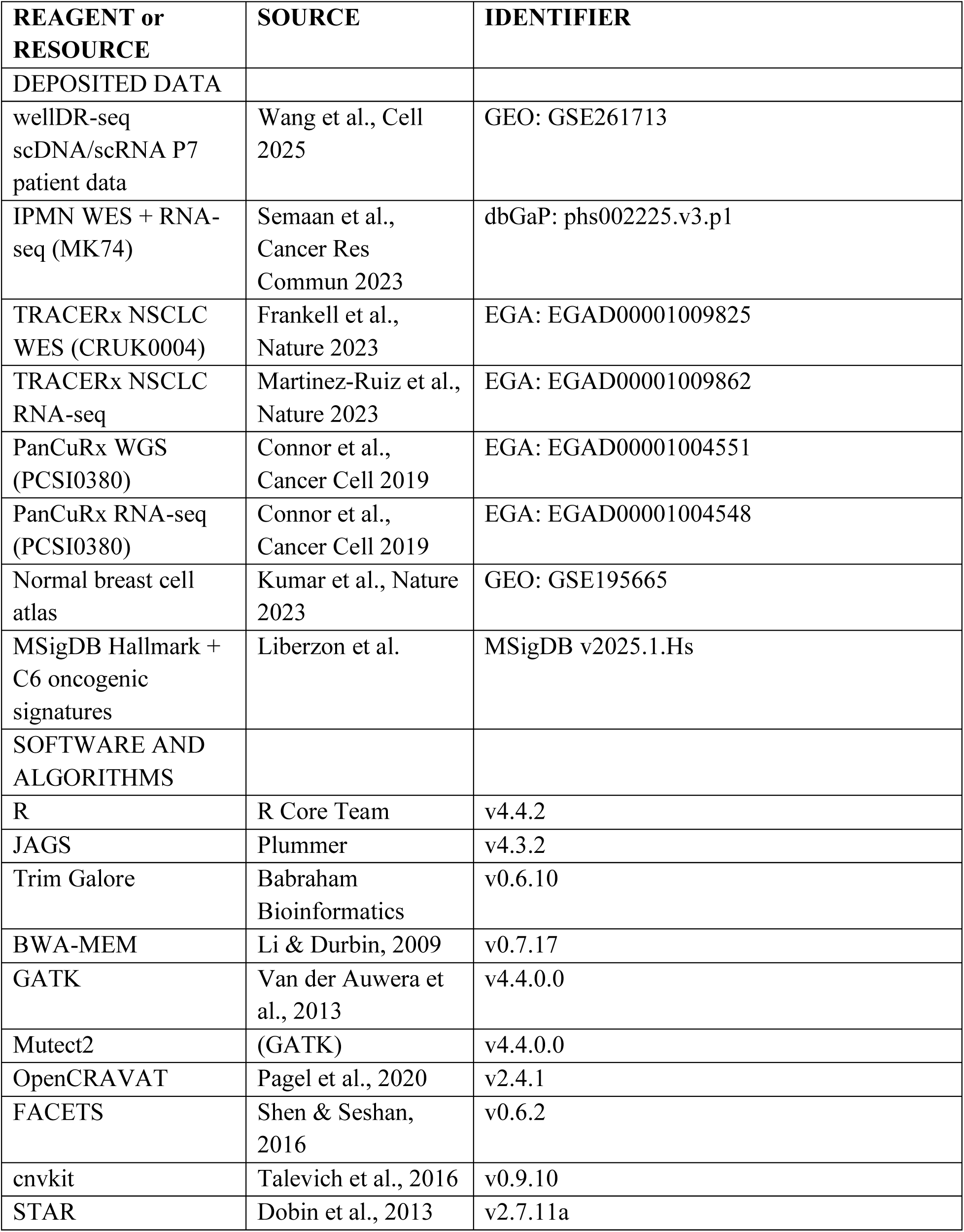
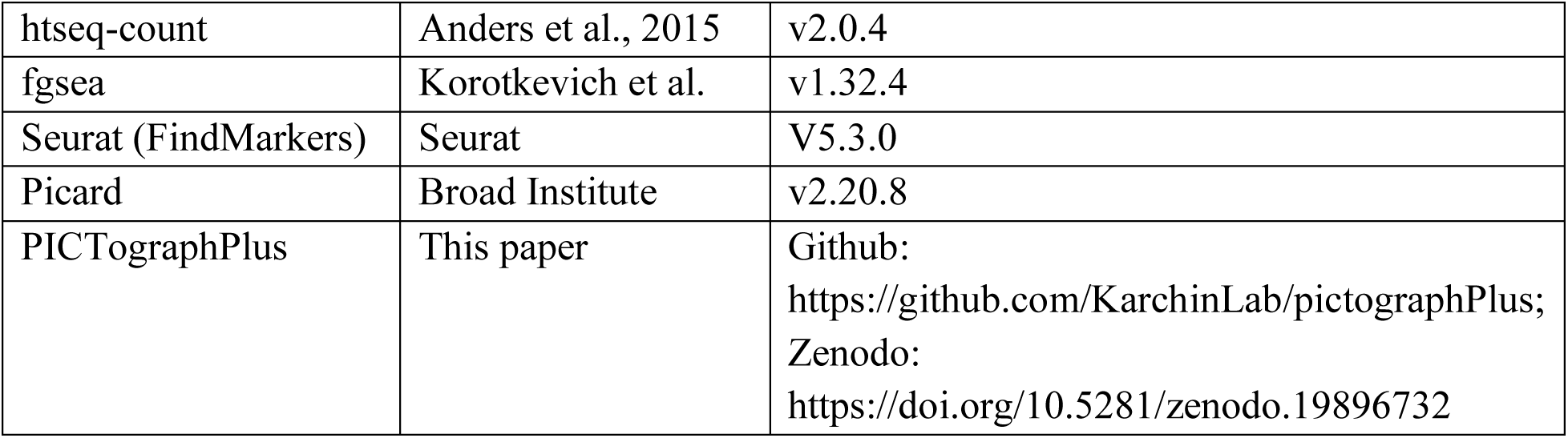

### Experimental model and subject/study participant details

This is a computational study using previously published, de-identified human tumor sequencing datasets. No new human subjects were enrolled. All primary datasets were generated under institutional review board approvals reported in the original studies: wellDR-seq (Wang et al., 2025) ^18^; TRACERx ^19–21^; NCI Cancer Moonshot Precancer Atlas Pilot^22^; and PanCuRx Translational Research Initiative^23^. All datasets were used in accordance with the respective data access agreements.

## Method details

### Tumor evolution reconstruction

Our tumor evolution reconstruction framework is designed to analyze multi-sample tumor genomic sequencing data from a single patient, either multi-region at one time point or longitudinally. Each sample is characterized by somatic single nucleotide variants (SNVs) and copy number alterations (CNAs). For SNVs, the inputs are the variant and total read counts; for CNAs, the input is the total copy number (TCN), defined as the sample-level copy number without purity adjustment. We adopt a partial infinite-sites assumption, under which each mutation can be acquired at most once but may be lost through a copy-number deletion.

To jointly estimate latent quantities of interest (mutation-cluster assignments, multiplicities, and mutation-cluster cellular fractions) from SNV and CNA data, we implement two hierarchical Bayesian generative models (Note S1). Model 1 estimates the cellular prevalence of CNAs, i.e. the CNA mutation cellular fraction (cncf), defined as the fraction of cells harboring a given copy-number alteration, using SNVs in copy-neutral regions together with all user-specified CNAs. Model 2 then refines parameter estimates by incorporating all SNVs and CNAs.

Both models use a Binomial likelihood for variant-read counts and a spike-and-slab prior on cluster cellular fractions to encourage sparsity of clone presence across samples. Segmented total copy number is modeled with a Normal likelihood, and priors on integer copy number (λ_icn = 2) and multiplicity (λ_m = 1) follow Poisson distributions with biologically motivated truncations. Models are implemented in JAGS (4.3.2) via R (v4.4.2) with 5,000 MCMC draws, thinning = 10, and 1,000 burn-in iterations. The optimal number of clusters can be selected using BIC or silhouette statistics^27,28^.

MCMC convergence was assessed for all three cases using split-chain Gelman-Rubin diagnostics. Because each JAGS model was run with a single chain, the posterior chain for each mutation cellular fraction (MCF) parameter was split at the midpoint into two halves, which were treated as independent chains; R-hat values close to 1.0 (threshold < 1.1) indicate that the two halves sampled the same distribution. Across all cases, 46 of 48 MCF parameters satisfied this criterion. Two parameters showed borderline values: mcf[2,3] in PCSI0380 (R-hat = 1.21, 97.5% CI upper bound 1.72) and mcf[6,2] in CRUK0004 (R-hat = 1.13, upper bound 1.34), both corresponding to clusters with low MCF in individual samples where the posterior is necessarily diffuse. All other parameters had R-hat ≤ 1.05, consistent with satisfactory convergence of the key model parameters.

Based on the posterior cluster mutation cellular fractions (MCFs), we enumerate candidate clone trees using a modified Gabow-Myers algorithm following pseudo-code from Popic et al.^24,25^.

Briefly, the algorithm recursively builds trees by adding edges only when parent-child pairs satisfy the lineage-precedence rule (the child MCF cannot exceed the parent MCF) and the Sum rule (the sum of children’s MCFs cannot exceed the parent MCF). This edge-adding restraint is what distinguishes the modified Gabow-Myers algorithm from the original. Small violations of each rule are tolerated up to a predefined threshold of 0.2. All trees that satisfy these constraints are returned. The resulting ensemble of trees is scored with a fitness function that penalizes rule violations,

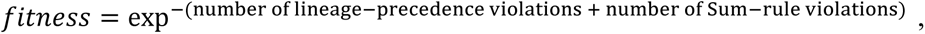

as in Niknafs et al. and Zheng et al.^3,16^.

### Clonal sample proportion estimation

Tumor clone proportions for each sample are derived from the inferred tree structure and cluster MCFs. For each clone in each sample, its “private” contribution is computed by subtracting the total MCF of its children (summed over all direct descendants in the tree) from the clone’s own MCF. Leaf clones therefore retain their full MCF, whereas internal clones contribute only the fraction not explained by their descendants. Any resulting negative values are truncated to zero, and the resulting subclone contributions are normalized so that subclone proportions sum to 1 within each sample. This procedure assumes that the input MCFs approximately respect both the lineage-precedence and Sum rules.

### Bulk RNA deconvolution: model specification

Let 𝑠 = number of samples, 𝑛 = number of genes, and 𝑘 = number of tumor clones plus the normal clone. Let 𝑌 be the 𝑠 by 𝑛 bulk gene expression matrix, 𝑋 the 𝑘 by 𝑛 matrix of clone-specific expression, Π the 𝑠 by 𝑘 matrix of clone proportions, 𝐿 the 𝑘 by 𝑘 normalized Laplacian matrix, 𝐸 be the set of tree edges, 𝑥_𝑘_ the expression vector for clone 𝑘, and for edge 𝑒 = (𝑝, 𝑐), let 𝑥_𝑐_ − 𝑥_𝑝_ denote the expression change across that edge. When a normal reference is included, an extra row Y_normal with Π_normal = [1, 0, …, 0] is appended, pinning the root/normal clone to a known reference.

All regularization models minimize a squared-residual data-fidelity term ||𝑌 − Π𝑋||^2^ plus a regularization penalty parameterized by a Laplacian L. Two choices of L were evaluated: a tree Laplacian (clonal phylogeny) and a star Laplacian (uniform smoothing to root). Based on topology-sensitivity analysis, we use the star Laplacian for all results reported.

We evaluated three deconvolution modes that differ in the normal reference appended to the bulk mixture: with-normal, which appends the matched within-dataset normal clone; with-external-normal, which appends an independent normal profile derived from a single-cell reference atlas (see below); and tumor-only, which includes no normal reference.

### Regularization models

We implemented and benchmarked seven regularization models. We summarize five primary objective classes here; details for each regularization model, as well as two additional implementations, are described in Note S2.

1. Plain Laplacian

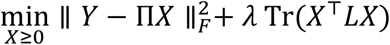
2. Adaptive Laplacian

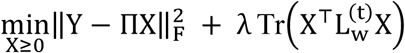
3. Fused edge-wise (fused_ew)

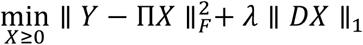

where

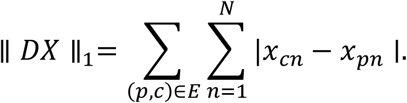
4. Elastic_net

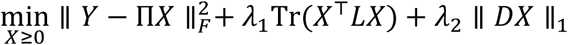
5. Tree_delta

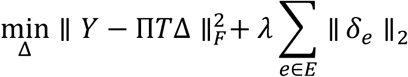

where 𝛿_𝑒_ ∈ ℝ^𝑁^ is the expression change associated with edge 𝑒.

### Benchmark dataset and pseudo-bulk simulation

We used patient P7 from the wellDR-seq ER+ breast cancer cohort, which simultaneously profiles DNA and RNA from individual single-cells^18^. scDNA identified one diploid normal clone (c1) and four tumor clones (c2, c4, c7, c11) that contained at least 30 cells, plus a rare subclone c13 (n = 16 cells) carrying only the chr16q loss, the founding copy-number event shared by all cancer subclones in P7. Because excluding c13 produces an artifactual polyclonal topology that contradicts the single clonal origin reported by Wang et al., we merged the 16 c13 cells into c11 (both sharing chr16q loss as their earliest event), yielding the trunk clone c11_c13 (n = 176 cells). This produces a monoclonal tree rooted at c1 with c11_c13 as a trunk that bifurcates into c2 (with child c7) and c4.

To construct a deeper benchmark tree with richer topology structure, we extended the four-clone tree by treating c7 and c4 as internal nodes and growing synthetic clones (c7_A, c7_B along the c7 lineage; c4_A, c4_B along the c4 lineage) via multiplicative perturbation of the parent’s pseudo-bulk profile using fold changes derived from five uniquely assigned Hallmark gene sets per synthetic clone (Table S6). All 20 assigned Hallmark gene sets are unique across synthetic clones. The resulting tree comprises 8 tumor clones with a maximum depth of 5 edges.

Ground-truth pathway enrichment for each tree edge was determined by fgseaMultilevel against the combined MSigDB Hallmark (H; 50 gene sets) and oncogenic signatures (C6) collections (201 sets after filtering for overlap with the expression matrix), with significance at BH-adjusted p < 0.05. The gene-level ranking fed to fgsea differs by edge type: for inherited edges (real scRNA-seq), Wilcoxon rank-sum statistics from FindMarkers (Seurat, min.pct = 0.10) between child and parent single-cells are used; for synthetic edges, pseudobulk log-fold-changes between the perturbed child profile and its parent are computed as log2((child+1)/(parent+1)), filtered to genes expressed in ≥10% of parent cells. edge_0_1 (root → c11_c13) yielded 49 significant pathways, edge_1_2 (c11_c13 → c2) yielded zero, and edges c11_c13 → c4 and c2 → c7 had low GT (excluded from F1).

For synthetic edges, two disjoint classes of genes are scaled ×5 in each child profile: (i) genes in the synthetic clone’s 5 assigned Hallmark sets, and (ii) the top 100 highest-expressed genes not belonging to any of the 20 assigned sets across all four synthetic clones. The second class is included to make the benchmark harder: their high parent-clone expression produces large absolute LFCs that drive enrichment of additional Hallmark and C6 sets overlapping those genes. As a result, GT significant-pathway counts substantially exceed 5: edge_3_4 = 53, edge_4_5 = 80, edge_6_7 = 38, edge_7_8 = 11. Ground truth is the full fgsea output, not the 5 assigned sets alone.

We generated 320 replicate pseudo-bulk datasets by mixing clone-level expression profiles using a stage-aware asymmetric Dirichlet sampling scheme. Each replicate drew clone proportions for each bulk sample independently from one of two evolutionary regimes (each with probability 0.5): early-stage, in which shallow clones (depth <= 2; c11_c13, c2, c4) dominate; and late-stage, in which deep synthetic clones dominate. Within each regime, one clone dominated in 70% of samples (Dirichlet alpha = 4.0; expected proportion ∼ 66%) and two co-dominated in the remaining 30% (alpha = 3.0 each). Non-dominant clones received background concentration alpha = 0.3. Simulations spanned four tumor purity levels (0.20, 0.40, 0.60, 0.80) and four numbers of tumor bulk samples (2, 3, 4, 5), with 20 random seeds per condition, yielding 320 replicates.

### External normal reference

To assess the effect of normal reference choice, we substituted the within-dataset normal clone (c1 cells, n = 297) with an independent external normal derived from a large-scale single-cell atlas of normal breast tissue (CellxGene Human Cell Atlas; 456,493 cells)^26^. We selected the top 4 most frequent cell types, spanning basal 22%, fibroblasts 46%, luminal epithelial 23%, vein endothelial 9%. Per-gene mean raw counts across these cells were scaled to the same reference cell count used for clone expression normalization (n_ref = 35), yielding a pseudo-bulk normal profile on the same scale as deconvolved clone expression. Gene space was restricted to the 14,647 genes shared between the P7 Seurat annotation (GENCODE locus-based identifiers) and the atlas HGNC symbol space; the 5,380 excluded genes are predominantly non-coding RNAs absent from tested gene sets.

### Baseline comparison

As a baseline, each clone was represented by the bulk sample with the highest estimated proportion of that clone. GSEA was then applied to differential expression between samples representing paired clones along each tree edge. This "max-proportion sample" baseline requires no explicit deconvolution and serves as the reference for evaluating whether the model-based approach adds value.

### Topology sensitivity and topology validity diagnostic

Topology sensitivity was evaluated under 17 tree topology variants in with-normal mode: the true tree, a star topology (all tumor clones equidistant from the root), five randomly permuted alternative topologies (alt x 5), five fixed-structure permutations (fixed x 5), and five linear chain topologies (linear x 5). For tumor-only and with-external-normal modes, only the true-tree and star tree topologies were evaluated. For each topology, the Laplacian matrix L was recomputed and deconvolution was performed across all 320 replicates at six regularization strengths.

Performance was evaluated by (i) expression recovery (Pearson r between deconvolved and true clone profiles, per clone per replicate) and (ii) edge-specific pathway recovery (GSEA F1 on synthetic and inherited edges). Topology advantage was quantified as the per-replicate difference (true tree minus mean across alternative topology families).

To investigate whether the true clonal phylogeny carries a measurable signal for clone expression similarity, a prerequisite for topology-based regularization to be useful, we computed pairwise expression similarity (Pearson r and cosine similarity) between all 8 tumor clone profiles and pairwise tree path lengths for all 17 topology variants. For each topology, we tested whether expression similarity decreases with tree distance using Spearman rank correlation and a Mantel permutation test (9,999 permutations, one-sided). This analysis provides an independent, upstream diagnostic of whether clone expression structure is consistent with a given phylogeny.

### Gene set enrichment analysis

Gene set enrichment analysis was performed to identify gene set-level changes between connected tumor clones. For each edge in the clone tree, we computed log2 fold changes by dividing the expression level of the child clone by that of its directly connected parent. Genes were ranked by these log ratios, and gene set enrichment was assessed using the fgsea package (version 1.32.4) with user-defined gene sets^29^. The combined MSigDB Hallmark (H; 50 gene sets) and oncogenic signatures (C6) collections (201 sets after filtering for overlap with the expression matrix) were used in the benchmarking. A customized gene set was used for case studies.

### DNA sequencing (WES/WGS)

For all cohorts, DNA sequencing reads were trimmed for quality and adapter removal using Trim Galore (version 0.6.10) and aligned to the human reference genome (hg38) using bwa mem (version 0.7.17) unless otherwise noted^30^. Alignments were processed following GATK (version 4.4.0.0) best-practices workflows for data pre-processing and somatic short variant discovery^31^. Somatic short variants were called using Mutect2 where WGS data were available and using the GATK somatic short variant pipeline for WES. SNVs were annotated using OpenCRAVAT (version 2.4.1)^32^, and only non-synonymous coding variants were retained for PICTographPlus.

### Copy-number calling

Copy number alterations (CNAs) were inferred from WES or WGS using FACETS (version 0.6.2), and, for IPMN, additionally using cnvkit (version 0.9.10)^33,34^. Total copy number (TCN) estimates were used as input to PICTographPlus without purity adjustment.

### RNA-seq processing

Unless otherwise specified, RNA-seq reads were trimmed with Trim Galore (version 0.6.10), aligned to hg38 using STAR (version 2.7.11a), and summarized to gene-level counts using htseq-count (version 2.0.4)^35,36^. When preprocessed RNA count matrices were provided by the original study authors, these were used directly.

### IPMN (Semaan et al.) data processing

Sequencing data for IPMN samples were obtained from Semaan et al., with whole-exome sequencing (WES) accessed via dbGaP (phs002225.v3.p1)^22^. SNVs were filtered to include only non-synonymous coding variants with minimum total read depth 30, variant read depth 10, and variant allele frequency (VAF) ≥ 0.05 in at least one sample. CNAs were identified using both cnvkit and FACETS as described above. Processed RNA read-count data were provided by the original study authors. PICTographPlus was run with default parameters, using mutation, copy-number, and germline heterozygous variant information as inputs.

### TRACERx NSCLC data processing

TRACERx NSCLC data were obtained from the European Genome-Phenome Archive (EGA) accessions EGAD00001009825 (WES) and EGAD00001009862 (RNA-seq)^19^. For downstream analysis, we retained only non-synonymous coding variants and applied cohort-specific filters: in at least one sample, the variant read count had to be ≥ 20, the total read count ≥ 100, and the VAF ≥ 0.05. CNAs and RNA-seq data were processed as described above. PICTographPlus was run with default parameters using both mutation and copy-number data as inputs.

### PanCuRx data processing

PanCuRx data were obtained from EGA accessions EGAD00001004548 (RNA-seq) and EGAD00001004551 (WGS)^23^. RNA-seq BAM files were already aligned to hg38; gene-level counts were generated with htseq-count as described above. WGS BAM files, initially aligned to hg19, were reprocessed to hg38 by reverting each BAM to unaligned reads with Picard RevertSam (version 2.20.8), converting to FASTQ with SamToFastq, realigning with bwa mem (version 0.7.17), and marking/removing duplicates with Picard MarkDuplicates. Mutations were included if, in at least one sample, the variant read count was ≥ 5, the total read count ≥ 40, and the VAF ≥ 0.10. CNAs were identified using FACETS and used to correct the cancer-cell fraction of mutations rather than as independent clonal features on the tree; PICTographPlus was run with default parameters in mutation-only mode.

## Quantification and statistical analysis

### Evaluation metrics

Expression recovery was quantified as the Pearson correlation between the deconvolved clone expression profile and the true synthetic profile, computed per clone per replicate and averaged across the 320 replicates. GSEA pathway recovery was quantified by comparing predicted significant pathways (adjusted p < 0.05) against ground-truth significant pathways per edge-replicate pair, using sensitivity (recall = TP / (TP + FN)), precision (TP / (TP + FP)), F1 score (harmonic mean of precision and sensitivity), and Matthews correlation coefficient (MCC = (TP×TN − FP×FN) / sqrt((TP+FP)(TP+FN)(TN+FP)(TN+FN))), with TP/FP/FN/TN computed relative to the combined MSigDB Hallmark (H; 50 gene sets) and oncogenic signatures (C6) collections (201 sets after filtering for overlap with the expression matrix). Primary GSEA metrics were computed on the four synthetic edges (controlled ground truth) and the informative inherited edge (edge_0_1, 49 ground-truth pathways). The null inherited edge (edge_1_2) and the low-GT inherited edges were excluded from F1 computation.

### Statistics

All hypothesis tests reported in this study are permutation-based (Mantel test, 9,999 permutations) or rank-based (Wilcoxon signed-rank for paired topology comparisons; Wilcoxon rank-sum for single-cell differential expression inputs to fgsea); these tests do not yield classical degrees of freedom, and we report exact permutation p values, test statistics (Mantel r, Wilcoxon W, Spearman ρ), and effect sizes (Pearson r). 95% confidence intervals on Pearson r values were computed by Fisher z-transformation of the per-replicate r values (n = 320 replicates) and back-transformation to the r scale; 95% confidence intervals on paired tree-vs-alternative differences were computed using the Hodges-Lehmann estimator (R wilcox.test with conf.int = TRUE).

Multiple comparisons across the four alternative-topology families (alt, fixed, linear, star) in the topology benchmark were Bonferroni-corrected.

### Declaration of generative AI and AI-assisted technologies in the manuscript preparation process

During the preparation of this work, the authors used Claude (Anthropic) for language editing and to check consistency of the manuscript text. The authors reviewed and edited the output as needed and take full responsibility for the content of the published article.

