## supplementary_file for "Reconstructing clone-resolved transcriptional programs from bulk tumor sequencing"

### Supplemental Information

- **Note S1**: Specification of the hierarchical Bayesian models
- **Note S2**: Seven regularization models for bulk RNA deconvolution
- **Figure S1**. True-tree expression recovery advantage
- **Figure S2**. GSEA topology comparison
- **Figure S3**. Pairwise expression similarity between tumor clones
- **Figure S4**. Expression recovery by deconvolution mode and clone type
- **Figure S5**. GSEA pathway recovery metrics for the root-adjacent edge
- **Figure S6**. GSEA pathway recovery metrics for the four synthetic edges
- **Figure S7.** Edge-wise gene set enrichment analyses along key evolutionary transitions in CRUK0004
- **Table S1.** Topology advantage of true tree compared to alternatives
- **Table S2.** Selected pairwise clone similarities and true-tree distances
- **Table S3.** Mantel / Spearman statistics across 17 topologies
- **Table S4.** Expression recovery at optimal regularization strength
- **Table S5.** GSEA pathway recovery on synthetic edges
- **Table S6.** Pathways altered for each synthetic edge

### Note S1: Specification of the hierarchical Bayesian models

#### Model intuition (overview)

PICTographPlus models clonal structure by clustering genomic events (SNVs and CNAs) into $K$ mutation clusters, each representing a latent clone whose prevalence varies across samples. For each cluster $k$ and sample $s$, the mutation cellular fraction $mcf_{ks}$ captures the fraction of tumor cells carrying events from that cluster, and is encouraged to be sparse across samples via a spike-and-slab prior (clones may be absent in some samples). We use two complementary likelihoods: (i) a Binomial model for variant/alternate allele counts derived from VAF/BAF, and (ii) a Gaussian model for segmentation-derived total copy number (TCN) for CNAs. We fit the model in two stages: Model 1 uses copy-neutral SNVs plus user-input CNAs to obtain stable initial estimates of cluster cellular fractions and CNA copy-number states; Model 2 then refines all parameters jointly using all SNVs and CNAs, additionally adjusting SNV expected VAFs for local CNA cellular fraction when an SNV lies within a CNA segment.

#### Notation and observed data

We analyze $I$ genomic events ("mutations"), including SNVs and CNAs, measured across $S$ bulk tumor samples from the same patient.

For each event $i \in \{1, \dots, I\}$ and sample $s \in \{1, \dots, S\}$, we observe:

- $h_i \in \{0,1\}$: event type indicator ($h_i = 0$ for SNV, $h_i = 1$ for CNA).
- $y_{is}$: variant read count for event $i$ in sample $s$.
- $n_{is}$: total read count for event $i$ in sample $s$.

For CNA events ($h_i = 1$), we additionally observe:

- $tcn_{is}$: continuous total copy number estimate from segmentation, interpreted as an average over a large chromosomal segment.

For CNAs, $y_{is}$ is derived from the B-allele frequency (BAF). The user may provide BAF directly, or PICTographPlus computes BAF from heterozygous SNPs for a given segment. Based on both TCN and BAF, PICTographPlus constructs pseudo-counts for $y_{is}$ and $n_{is}$.

##### Latent variables and parameters

- $VAF_{is}$: true variant allele frequency for event $i$ in sample $s$.
- $K$: number of mutation clusters.
- $Z_i \in \{1, \dots, K\}$: cluster assignment of event $i$.
- $\pi = (\pi_1, \dots, \pi_K)$: mixture weights over clusters.
- $mcf_{ks} \in [0,1]$: mutation cellular fraction for cluster $k$ in sample $s$.
- $cncf_{is} \in [0,1]$: copy-number cellular fraction (fraction of tumor cells in sample $s$ carrying a CNA segment overlapping event $i$); used to adjust SNV VAFs when an SNV lies in a CNA region.
- $icn_i$: integer copy number state associated with event $i$ (diploid baseline $icn_i = 2$ for copy-neutral).

- $m_i$: multiplicity (number of mutated copies) for event $i$.
- $M_i$: major integer copy number, derived as $M_i = max\ (m_i, icn_i - m_i)$ (used in Model 2).

**Mixture model for clonal structure**

Cluster assignments follow a categorical mixture model:

$Z_i|\ \pi \sim Categorical(\pi_1, \dots, \pi_K)$, where $\pi \sim Dirichlet(1_1, \dots, 1_K)$

**Spike-and-slab prior on cluster cellular fractions**

To encourage sparsity of clone presence across samples, we place a spike-and-slab prior on $mcf_{ks}$:

$\eta_{ks} \sim Bernoulli(1-\theta)$, $\theta \sim Beta(5,2)$

$$mcf_{ks}|\ \eta_{ks} \sim \begin{cases} 0, \ \eta_{ks} = 0 \\ Beta(1,1), \ \eta_{ks} = 1 \end{cases}$$

**Two-stage modeling approach**

We fit the model in two stages. Model 1 estimates cluster-level cellular fractions for CNA events (interpreted as CNA cellular fractions) using (i) SNVs in copy-neutral regions and (ii) user-input CNAs. Model 2 refines all parameters jointly using all SNVs and CNAs, and uses the CNA cellular fraction estimated from Model 1 to adjust SNV VAF expectations in CNA-overlapping regions. In cases where the user has already obtained CNA cellular fraction, Model 2 can be applied directly to SNVs.

#### Model 1 (initialization using copy-neutral SNVs + CNAs)

**Likelihood for VAF-derived counts**

For all events $i$ and sample $s$:

$y_{is}|\ n_{is}, VAF_{is} \sim \ Binomial(n_{is}, VAF_{is})$,

Let $z = Z_i$. $VAF_{is}$ is calculated by:

$$VAF_{is}|Z_i = z, h_i, mcf_{zs}, tcn_{is}, m_i = \begin{cases} \dfrac{mcf_{zs}}{2}, \quad h_i = 0\ (SNV, copy\ neutral) \\ \dfrac{mcf_{zs} * m_i + (1 - mcf_{zs})}{tcn_{is}}, h_i = 1\ (CNA, BAF\ derived) \end{cases}$$

**Likelihood for segmented total copy number (CNA only)**

For CNA events ($h_i = 1$), we model segmented total copy number as approximately normal:

$$tcn_{is}|Z_i = z, icn_i, mcf_{zs} \sim Normal(icn_i * mcf_{zs} + 2 * (1 - mcf_{zs}), \sigma_i)$$

This is motivated by $tcn_{is}$ being an average over many loci within a segment (law of large numbers). We place a half-normal prior on the standard deviation, $\sigma_i \sim Half - Normal(0,1)$ (i.e., a $Normal(0,1)$ distribution truncated to $\sigma_i > 0$), in the implementation.

**Priors for integer copy number and multiplicity**

We use:

$$icn_i | h_i = \begin{cases} 2, h_i = 0 \\ Poisson(\lambda_{icn}), h_i = 1 \end{cases}$$

$$m_i | icn_i, h_i = \begin{cases} 1, \ h_i = 0 \\ \min(Poisson(\lambda_m), icn_i), \quad h_i = 1 \end{cases}$$

In our implementation, we set $\lambda_{icn} = 2$ and $\lambda_m = 1$, with truncation to ensure $1 \le m_i \le icn_i$.

**Outputs carried to Model 2**

From model 1, we retain:

- $mcf_{ks}$ for CNA clusters, which are used as CNA cellular fraction estimates $cncf_{is}$ for CNA-overlapping adjustments in Model 2.
- $icn_i$ and $m_i$, from which we compute $M_i = max\ (m_i, icn_i - m_i)$ for use in Model 2.

#### Model 2 (joint refinement using all SNVs + CNAs)

**Likelihood for VAF-derived counts**

As in Model 1:

$y_{is} |\ n_{is}, VAF_{is} \sim Binomial(n_{is}, VAF_{is})$,

**VAF expectation with CNA-overlap adjustment for SNVs**

Let $z = Z_i$. In Model 2, the expected VAF is:

$$VAF_{is} | Z_i = z, h_i, mcf_{zs}, icn_i, m_i, cncf_{is} = \begin{cases} \dfrac{mcf_{zs} + (m_i - 1)cncf_{is}}{icn_i * cncf_{is} + 2(1 - cncf_{is})}, & h_i = 0\ (SNV) \\ \dfrac{mcf_{zs} * m_i + (1 - mcf_{zs})}{icn_i * mcf_{zs} + 2(1 - mcf_{zs})}, & h_i = 1\ (CNA) \end{cases}$$

Here, $cncf_{is}$is treated as fixed/initialized from Model 1 for CNA-overlapping regions.

**Multiplicity prior in Model 2**

We constrain multiplicity using the major copy number $M_i$:

$m_i | icn_i, M_i \sim Categorical(1, M_i, icn_i - M_i)$,

i.e., $m_i$ is restricted to these three biologically plausible values (with equal prior mass unless otherwise specified).

**Cluster priors**

As in Model 1:

$Z_i |\ \pi \sim Categorical(\pi_1, \ldots, \pi_K),\ \pi \sim Dirichlet(1_1, \ldots, 1_K)$,

and $mcf_{ks}$ follows the same spike-and-slab prior defined above.

### Note S2: Seven regularization models for bulk RNA deconvolution

Note S2 specifies all seven regularization models. The five highlighted in the main text are the plain Laplacian, adaptive Laplacian, fused edge-wise, elastic-net tree, and tree-delta models (models 1, 2, 5, 6, and 7 below); the adaptive Laplacian v2 (model 3) and plain debiased (model 4) are close variants of the adaptive and plain Laplacian models, respectively, included here for completeness.

Let:

- $Y \in \mathbb{R}^{S\times N}$ be the observed bulk expression matrix for $S$ samples and $N$ genes,
- $\Pi \in \mathbb{R}^{S\times K}$ be the clone proportion matrix,
- $X \in \mathbb{R}^{K\times N}$ be the unknown clone-level expression matrix,
- $L \in \mathbb{R}^{K\times K}$ be the graph Laplacian of the clone tree,
- $E$ be the set of tree edges,
- $x_k \in \mathbb{R}^N$ denote the expression vector for clone $k$,

For edge $e = (p, c)$, where p is the parent clone and c is the child clone, let $x_c - x_p$ denote the expression change across that edge.

**Laplacian convention**

Throughout Note S2, every occurrence of L (the plain Laplacian (Models 1, 4), the weighted Laplacian L_w in the IRLS updates (Models 2, 3), and the Laplacian term in the elastic-net tree model (Model 6)) refers to the symmetric (spectrally) normalized Laplacian,

$$L = D^{-1/2}(D - A)D^{-1/2}$$

where A is the adjacency matrix of the clone tree and D is the corresponding degree matrix. For Models 2 and 3 the normalization is recomputed each IRLS iteration from the updated edge weights $w_{pc}^{(t)}$. The normalization rescales each clone's contribution by $\frac{1}{\sqrt{d_k}}$, so that penalty strength is comparable across topologies of unequal root degree (a star tree, for example, has root degree K − 1 while every leaf has degree 1) and the linear systems remain well-conditioned. Consequently, the identity $\mathrm{Tr}(X^\top LX) = \sum_{(p,c)\in E} \left\| x_c - x_p \right\|_2^2$ that holds for the unnormalized Laplacian no longer applies; instead, the penalty takes the degree-rescaled form $\mathrm{Tr}(X^\top LX) = \sum_{(p,c)\in E} \left\| \frac{x_c}{\sqrt{d_c}} - \frac{x_p}{\sqrt{d_p}} \right\|_2^2$. Models 5 (fused edge-wise) and 7 (tree delta) do not use the Laplacian and are unaffected.

**1. Plain Laplacian model**

The plain Laplacian model solves

$$\hat{X} = \arg\min_{X\geq 0}\{\|Y - \Pi X\|_F^2 + \lambda \mathrm{Tr}(X^\top LX)\}$$

The Laplacian penalty encourages expression profiles of adjacent clones to be similar; under the normalization defined above, it penalizes degree-rescaled differences along each edge. For numerical implementation, we first compute the unconstrained ridge-stabilized solution

$$\tilde{X} = (\Pi^T \Pi + \lambda L + \varepsilon I)^{-1} \Pi^T Y$$

where $\varepsilon I$ is a small diagonal stabilizer. Nonnegativity is then enforced by elementwise projection,

$$\hat{X} = \max(\tilde{X}, 0).$$

This projection is used as a computational approximation to the nonnegative penalized problem.

**2. Adaptive Laplacian**

The adaptive Laplacian model replaces the uniform edge penalty with edge-specific weights:

$$X = \arg\min_{X \geq 0} \{ \|Y - \Pi X\|_F^2 + \lambda \operatorname{Tr}(X^\top L_w^{(t)} X) \}$$

Where $L_w^{(t)} = D_w^{(t)-1/2} \left( D_w^{(t)} - A_w^{(t)} \right) D_w^{(t)-1/2}$, with edge weights $w_{pc}^{(t)}$ updated each IRLS iteration as defined below:

$$w_{(pc)}^{(t+1)} = \frac{1}{\left\| x_c^{(t)} - x_p^{(t)} \right\|_2 + \epsilon}$$

where $\epsilon > 0$ is a small constant for numerical stability.

**3. Adaptive Laplacian v2 (adaptive_v2)**

This model has the same objective as the adaptive Laplacian,

$$X = \arg\min_{X \geq 0} \{ \|Y - \Pi X\|_F^2 + \lambda \operatorname{Tr}(X^\top L_w^{(t)} X) \}$$

with the same IRLS update

$$w_{pc}^{(t+1)} = \frac{1}{\| x_c^{(t)} - x_p^{(t)} \|_2 + \epsilon}.$$

The difference is only in initialization. Instead of starting from uniform weights, the initial estimate is taken from the unregularized fit

$$X^{(0)} = \arg\min_{X \geq 0} \| Y - \Pi X \|_F^2,$$

and the first set of weights is computed from $X^{(0)}$.

**4. Plain debiased Laplacian model**

First, compute the penalized plain Laplacian estimate

$$\hat{X}_{\text{pen}} = \arg\min_{X \geq 0} \{ \| Y - \Pi X \|_F^2 + \lambda \operatorname{Tr}(X^\top L X) \}.$$

Then apply the first-order debiasing correction

$$\hat{X}_{\text{deb}} = \hat{X}_{\text{pen}} + \lambda (\Pi^\top \Pi + \varepsilon I)^{-1} L \hat{X}_{\text{pen}}.$$

If a truncated SVD is used for numerical stability, let

$$\Pi^\top \Pi = U D U^\top,$$

and retain only singular values $d_j$ satisfying $d_j > \tau \max_\ell d_\ell$, with $\tau = 0.01$. Then

$$(\Pi^\top \Pi)_\tau^\dagger = U_\tau D_\tau^{-1} U_\tau^\top,$$

and the correction becomes

$$\hat{X}_{\text{deb}} = \hat{X}_{\text{pen}} + \lambda (\Pi^\top \Pi)_\tau^\dagger L \hat{X}_{\text{pen}}.$$

**5. Fused edge-wise model**

The fused edge-wise model encourages sparse gene-level expression changes along tree edges. Let $D \in \mathbb{R}^{|E| \times K}$ be the signed edge-incidence matrix of the clone tree. The model solves

$$\hat{X} = \arg \min_{X \geq 0} \{\| Y - \Pi X \|_F^2 + \lambda \| DX \|_1\},$$

where

$$\| DX \|_1 = \sum_{(p,c) \in E} \sum_{n=1}^{N} |x_{cn} - x_{pn}|.$$

Unlike the Laplacian penalty, which applies an $L_2$-type smoothness penalty across all genes, the fused edge-wise penalty uses an $L_1$-type penalty on individual gene-level changes. This encourages many edge-specific gene expression changes to be exactly zero while allowing a subset of genes to change sharply along specific clone-tree edges.

**6. Laplacian-fused elastic-net tree model**

The elastic net tree model combines Laplacian smoothing and edge-wise sparsity:

$$\hat{X} = \arg \min_{X \geq 0} \{\| Y - \Pi X \|_F^2 + \lambda_1 \operatorname{Tr}(X^\top L X) + \lambda_2 \| DX \|_1\}.$$

We refer to this as an elastic-net tree penalty because it combines an $L_2$-type Laplacian penalty with an $L_1$-type fused edge penalty. The Laplacian term promotes global smoothness of expression across the tree, whereas the fused term encourages sparse, edge-specific gene expression changes.

**7. Tree Delta**

In the tree-delta model, clone expression is parameterized as cumulative edge changes from the root. Let $\mu \in \mathbb{R}^N$ be the root expression vector and let $\delta_e \in \mathbb{R}^N$ be the expression change associated with edge $e$. Then for clone $k$,

$$x_k = \mu + \sum_{e \in \text{path}(r \to k)} \delta_e.$$

Stacking these terms, define $\Delta \in \mathbb{R}^{(1+|E|) \times N}$ with first row $\mu$ and remaining rows $\delta_e$, and let $T \in \{0,1\}^{K \times (1+|E|)}$ be the path matrix such that

$$X = T\Delta.$$

The optimization problem is

$$\hat{\Delta} = \arg \min_{\Delta} \left\{ \| Y - \Pi T \Delta \|_F^2 + \lambda \sum_{e \in E} \| \delta_e \|_2 \right\}.$$

Then the estimated clone expression matrix is

$$\hat{X} = T\widehat{\Delta}.$$

The penalty is group-sparse at the edge level, encouraging some entire edge-delta vectors to be exactly zero.

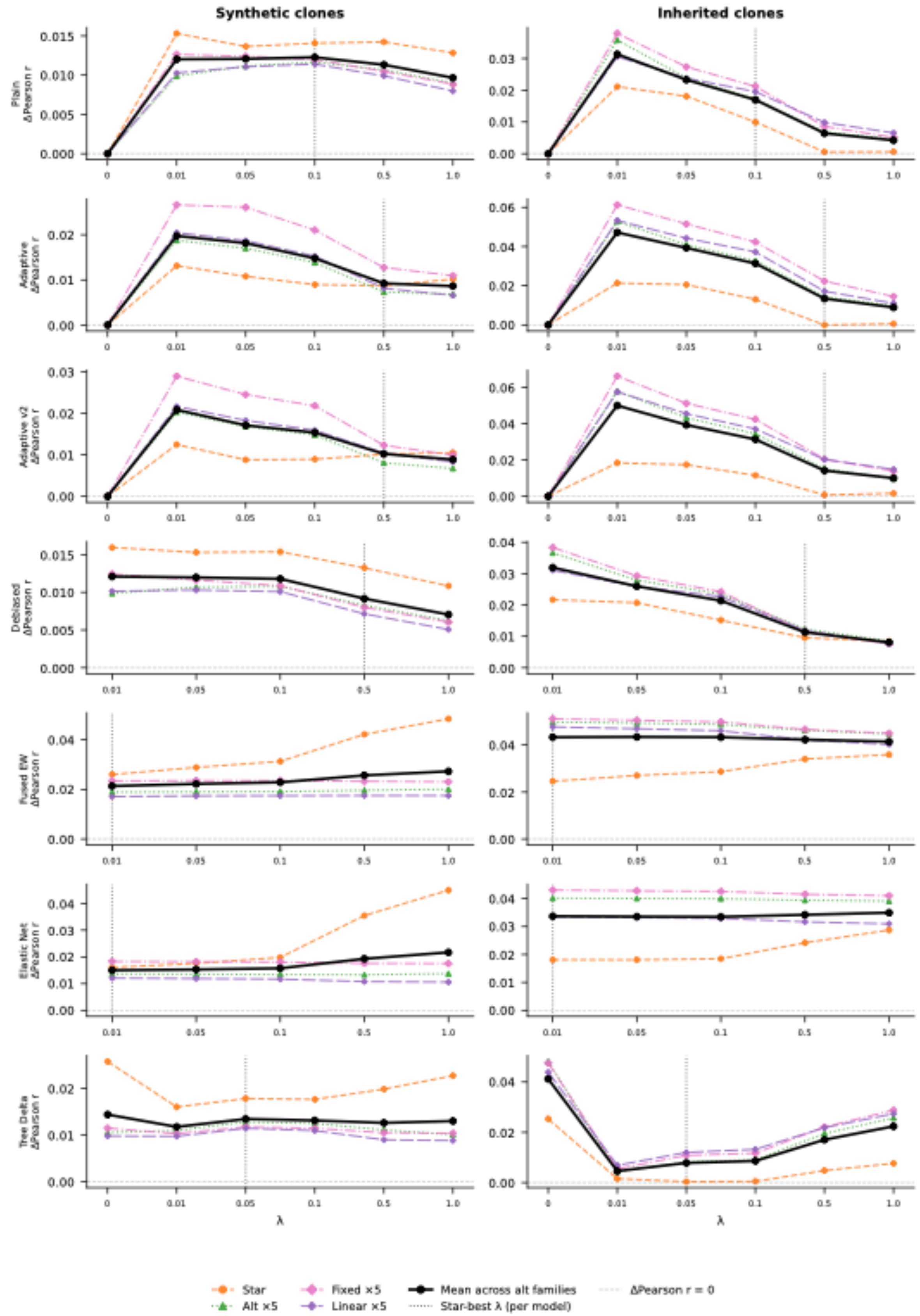

Synthetic clones
Inherited clones
Plain ΔPearson r
Adaptive ΔPearson r
Adaptive v2 ΔPearson r
Debiased ΔPearson r
Fused EW ΔPearson r
Elastic Net ΔPearson r
Tree Delta ΔPearson r
λ
Star
Alt ×5
Fixed ×5
Linear ×5
Mean across alt families
Star-best λ (per model)
ΔPearson r = 0

**Figure S1**. True-tree expression recovery advantage. Rows = 7 regularization models; left = 4 synthetic clones; right = 4 inherited clones. Each panel shows ΔPearson r = r(true tree) - r(alternative family), averaged over 320 replicates in with-normal. Line colors: star (orange), alt×5 (green), fixed×5 (pink), linear×5 (purple). Thick black = mean across the four alternative families. Vertical dotted line = star-tree-best λ per model. All Δr values are positive, confirming a small but consistent true-tree advantage in per-clone expression recovery over every alternative family.

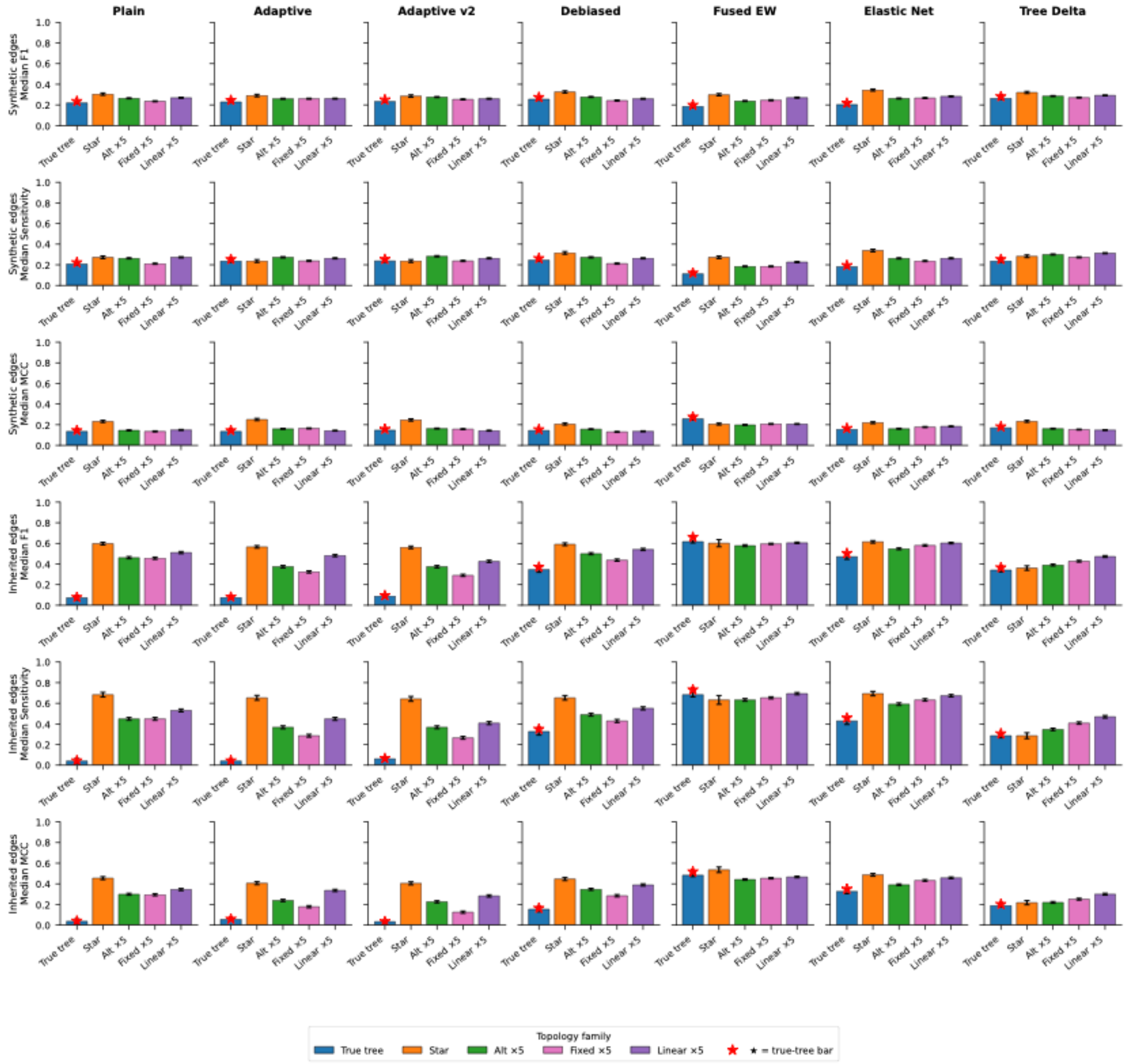

**Figure S2.** GSEA topology comparison. For each model (columns), median GSEA Sensitivity, F1, and MCC across all five topology families (the true tree and four alternative families: star, alt×5, fixed×5, linear×5). Top three rows: synthetic edges (depths 3-5). Bottom three rows: inherited edges (depths 1-2). Red star marks the true-tree bar. With-normal only. Across models and metrics, the true tree does not consistently outperform the alternatives in pathway recovery: the small expression-recovery advantage seen in Figure S1 does not translate into better pathway-level detection, consistent with the weak topology signal.

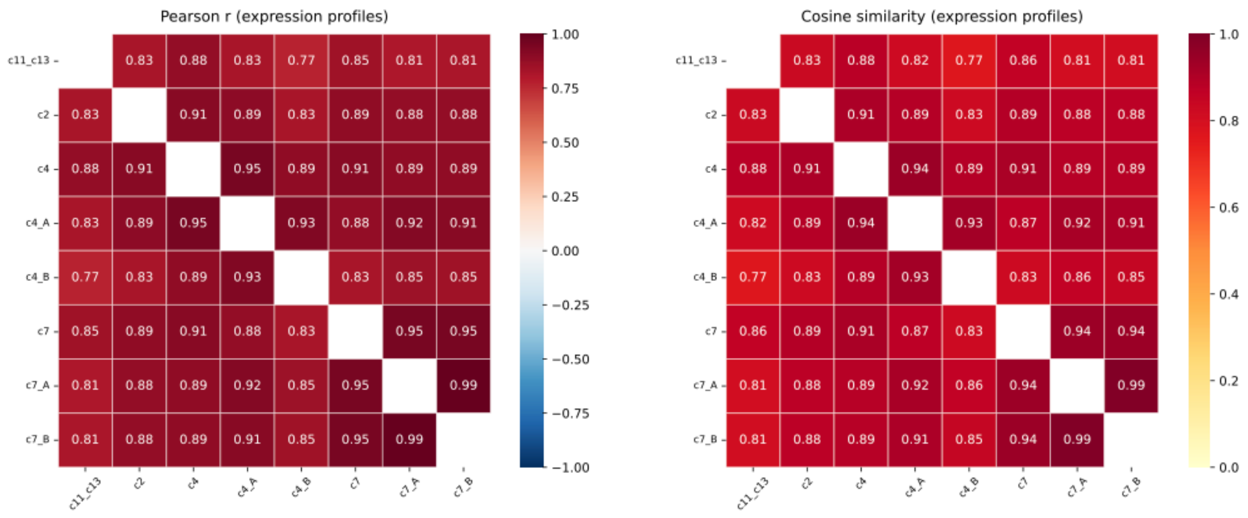


**Figure S3.** Pairwise expression similarity between tumor clones. Left panel: Pearson r similarity matrix. Right panel: cosine similarity. Symmetric 8×8 matrices on pseudo-bulk profiles of all eight tumor clones from scRNA-seq, restricted to genes with mean count ≥ 1.

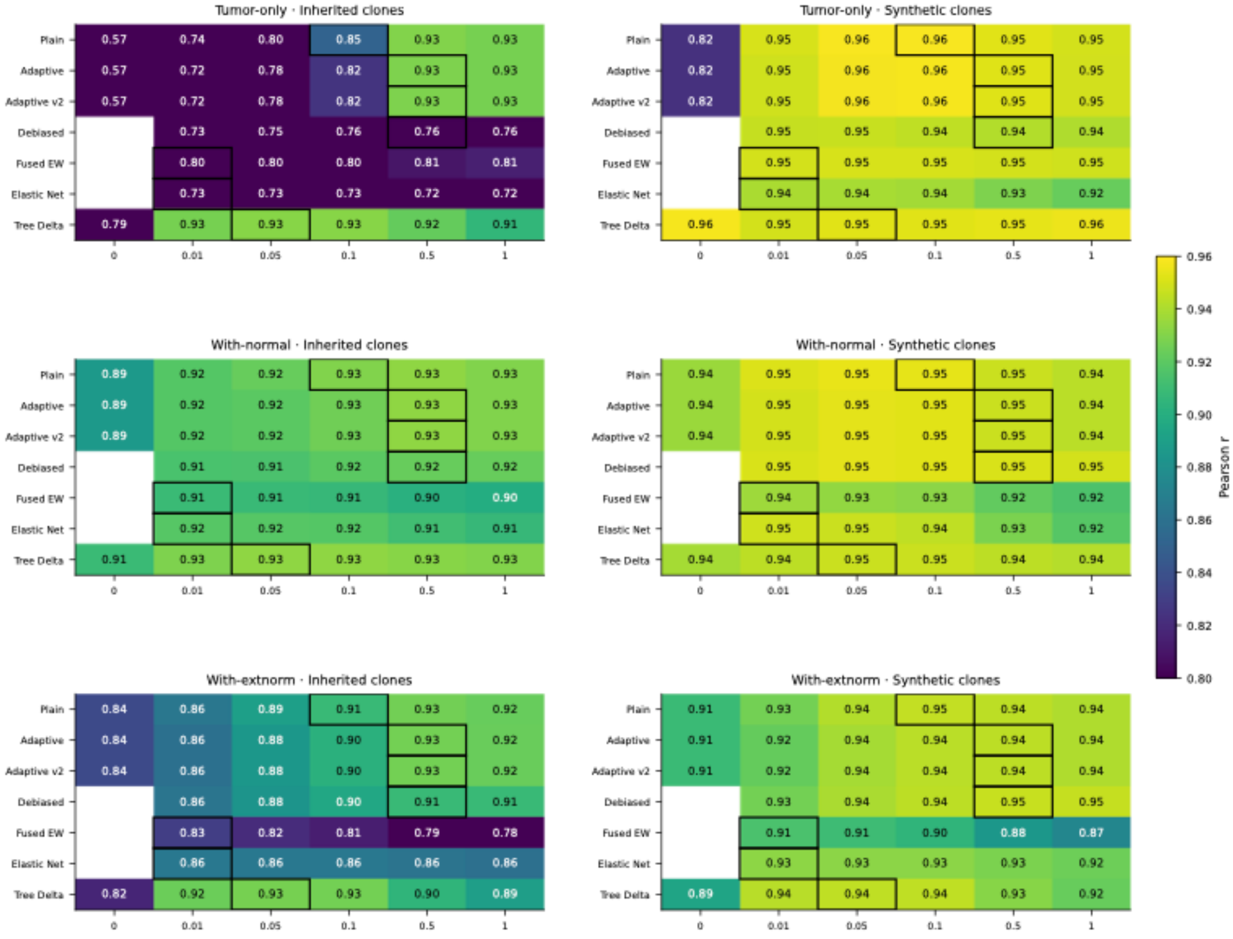


**Figure S4.** Expression recovery by deconvolution mode and clone type (star topology). Each panel shows mean Pearson correlation between deconvolved clone expression profiles and true synthetic profiles, averaged across all 320 replicates on the star topology. Rows correspond to the three deconvolution modes: tumor-only (no reference normal sample), with-normal (one matched normal sample included), and with-external-normal (population-average normal used). Left column: inherited (real single-cell) clones; right column: synthetic clones carrying Hallmark gene set perturbations. Black borders indicate the star-tree-best λ per model (a topology-agnostic selection criterion).

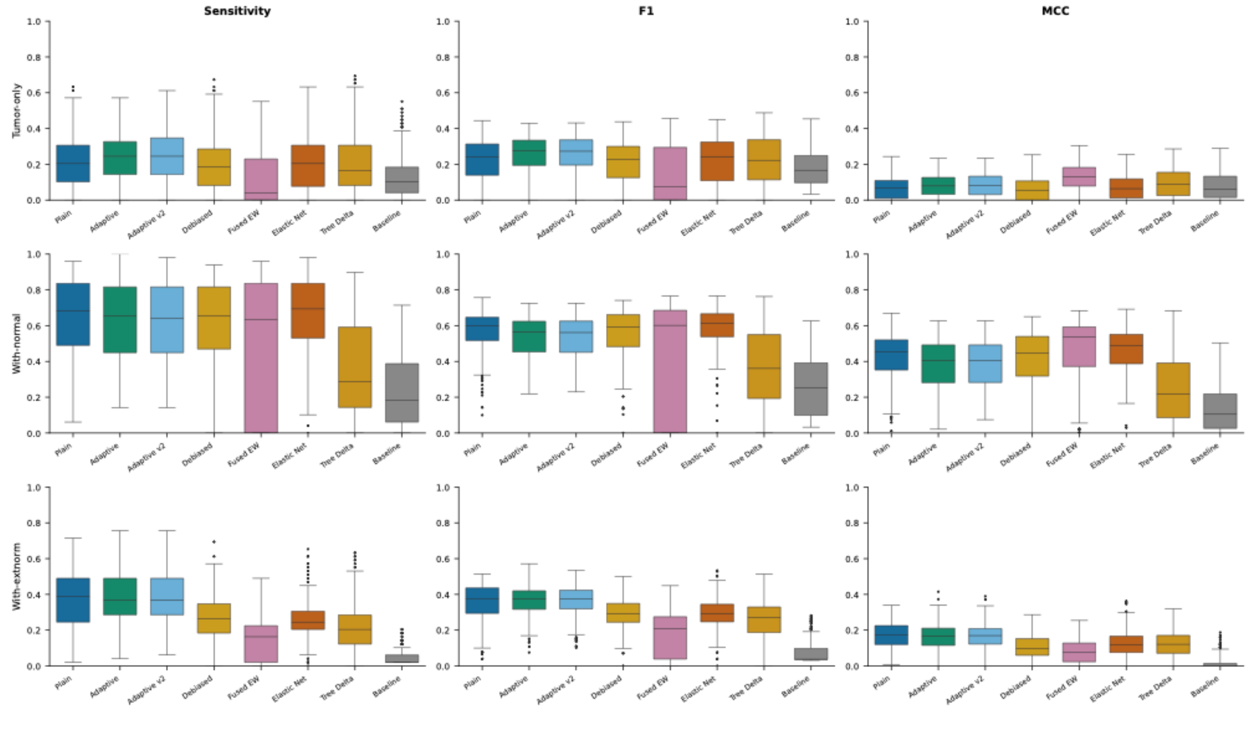


**Figure S5.** GSEA pathway recovery metrics (Sensitivity, F1 score, Matthews correlation coefficient) for the root-adjacent edge across three deconvolution modes (tumor-only, with-normal, with-external-normal). Each panel is a boxplot comparing the seven regularization models at their star-tree-best λ values across 320 replicates. The normal reference sample provides a substantial and consistent benefit across six of the seven models, with median F1 increasing approximately 2.0-2.6 fold from tumor-only (0.22-0.28) to with-normal mode (0.56-0.62). Tree Delta is an exception, achieving lower F1 (0.36) in with-normal mode due to the tree-structured penalty shrinking the trunk log fold-change under the uninformative star prior. The benefit reflects the mechanistic role of anchoring: the normal sample pins the root clone expression to a known reference, disambiguating the log fold-change on this root-to-trunk edge.

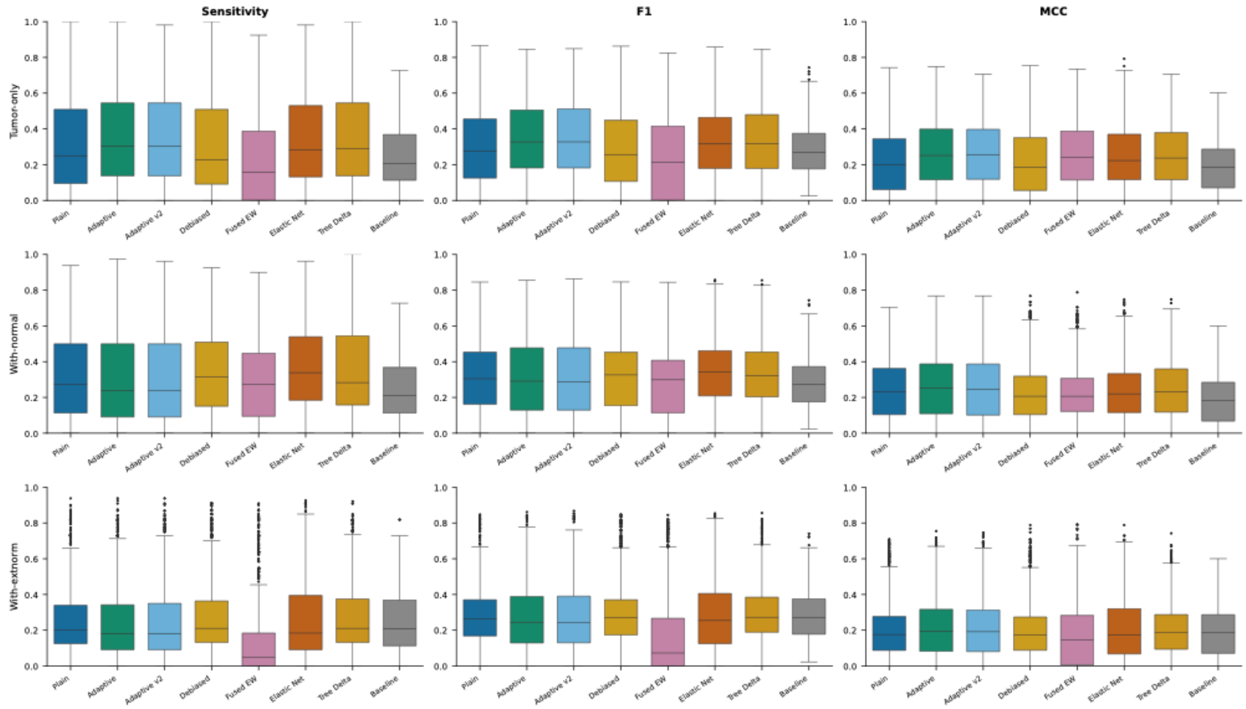


**Figure S6.** GSEA pathway recovery metrics (Sensitivity, F1 score, Matthews correlation coefficient) for the four synthetic edges at depths 3-5, stratified by deconvolution mode (tumor-only, with-normal, with-external-normal). Each panel is a boxplot comparing the seven regularization models at their star-tree-best λ values across 320 replicates per edge. Unlike the root-connected edge, the normal reference provides minimal additional benefit at the star-best λ for these tumor-to-tumor edges: median F1 values are comparable between tumor-only and with-normal modes. This pattern reflects the structure of the deconvolution problem: on edges between two tumor clones, the GSEA signal is the log fold-change between child and parent expression, a relative quantity that does not directly depend on the absolute root-scale anchor. The normal sample constrains the tumor-root pair but not the relative differences among tumor clones, so it only provides direct benefit to the root-to-trunk edge.

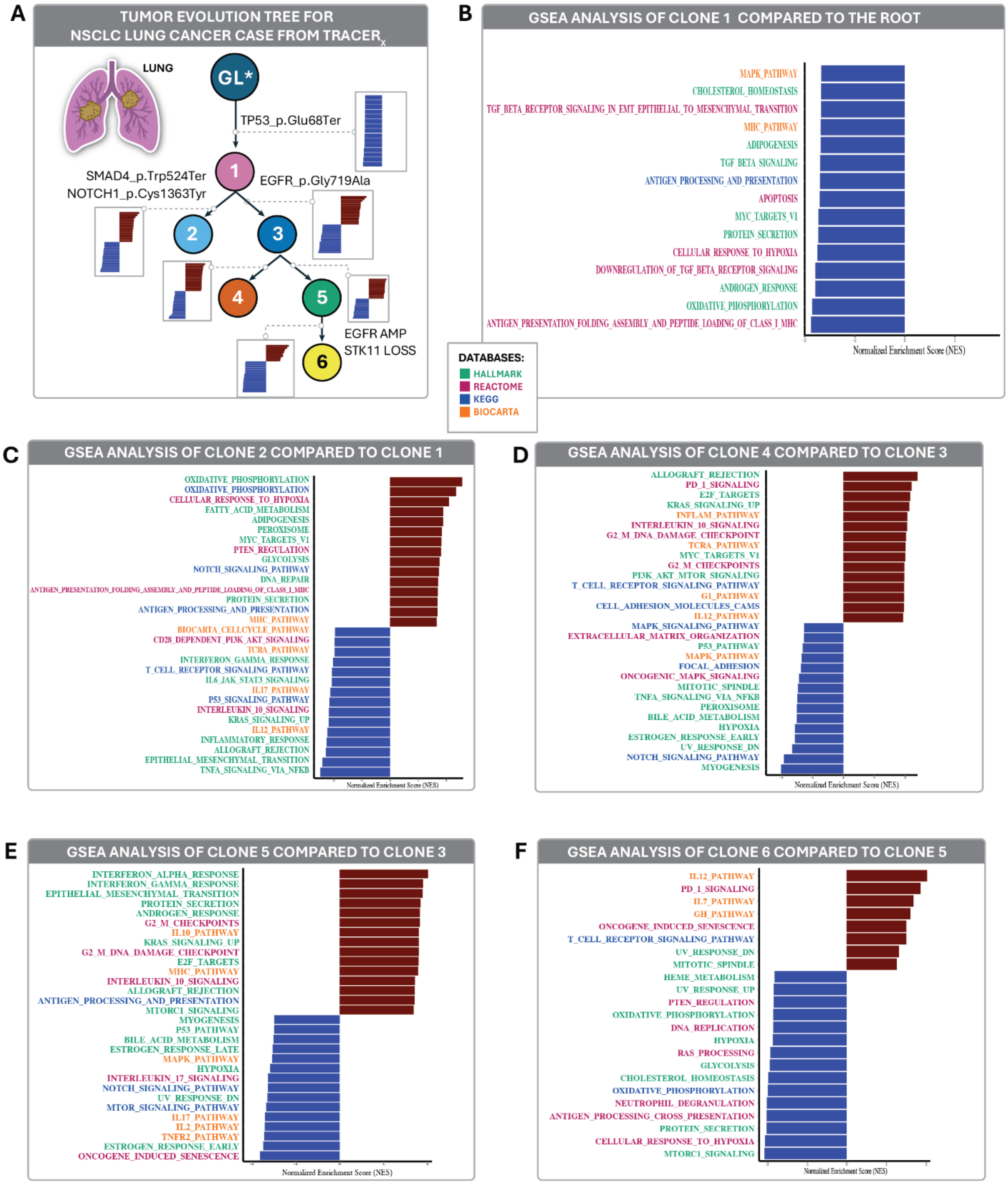


**Figure S7. Edge-wise gene set enrichment analyses along key evolutionary transitions in CRUK0004.** A) Clonal tree with truncal TP53 p.Glu68Ter (nonsense), SMAD4/NOTCH1 on Clone 2 branch, EGFR on Clone 3, terminal EGFR amp + STK11 loss on Clone 6. B) GSEA of the GL to Clone 1 transition, showing broad suppression of oxidative phosphorylation, MHC class I antigen presentation, MYC targets, TGF-β, hypoxia, apoptosis, cholesterol homeostasis, and MAPK signaling relative to normal lung, with no significant positive enrichments. C) GSEA of the Clone 1 to Clone 2 transition: upregulation of oxidative phosphorylation, glycolysis, fatty acid metabolism, peroxisome, adipogenesis, hypoxia response, MYC targets, and DNA repair; suppression of TNFα-NFκB, EMT, inflammatory response, IL6-JAK-STAT3, KRAS-up, and interferon-γ. D) GSEA of the Clone 3 to Clone 4 transition: upregulation of immune and proliferative programs including allograft rejection, PD-1, T-cell receptor, IL-10/IL-12, E2F targets, G2/M checkpoints, MYC targets, and PI3K-AKT–mTOR. E) GSEA of the Clone 3 to Clone 5 transition: upregulation of interferon-α/γ, EMT, G2/M, E2F targets, antigen presentation, KRAS-up, and mTORC1; suppression of oncogene-induced senescence, mTOR (KEGG), estrogen response, TNFR2, IL-2/IL-17, and p53 pathway activity. F) GSEA of the terminal Clone 5 to Clone 6 transition: widespread negative enrichment of metabolic and stress-response pathways (mTORC1, oxidative phosphorylation, glycolysis, hypoxia, cholesterol homeostasis, heme metabolism, DNA replication, neutrophil degranulation, antigen cross-presentation) together with modest upregulation of PD-1 signaling, IL-12, IL-7, and T-cell receptor signaling, consistent with metabolic and immune reprogramming in the metastatic clone following systemic therapy. Enrichment scores were computed using Hallmark, Reactome, KEGG, and BIOCARTA gene sets; database source is indicated by bar color.

Table S1**.** Topology advantage (true tree - mean alternative) at λ = 0.1: Hodges-Lehmann estimator with 95% confidence intervals from paired Wilcoxon signed-rank test (320 replicates, synthetic clones, with-normal mode, Bonferroni-corrected across four alternative-topology families).

| Model | Mean Δr | H-L estimate | 95% CI | Paired Wilcoxon p |
|---|---|---|---|---|
| Plain | +0.0118 | +0.0131 | [+0.0120, +0.0140] | < 0.001 |
| Adaptive | +0.0162 | +0.0180 | [+0.0162, +0.0199] | < 0.001 |
| Adaptive_v2 | +0.0170 | +0.0192 | [+0.0175, +0.0210] | < 0.001 |
| Plain_debiased | +0.0109 | +0.0120 | [+0.0110, +0.0130] | < 0.001 |
| Fused_ew | +0.0207 | +0.0236 | [+0.0210, +0.0265] | < 0.001 |
| Elastic_net | +0.0147 | +0.0167 | [+0.0149, +0.0187] | < 0.001 |
| Tree_delta | +0.0120 | +0.0124 | [+0.0116, +0.0132] | < 0.001 |

Table S2. Selected pairwise clone similarities and true-tree distances (Pearson r).

| Clone pair | Pearson r | True tree distance |
|---|---|---|
| c4 - c4_A (parent-child) | 0.948 | 1 |
| c7 - c7_A (parent-child) | 0.946 | 1 |
| c2 - c4 (siblings) | 0.907 | 2 |
| c4 - c7 (cross-branch) | 0.907 | 3 |
| c4_A - c7_A (cross-branch) | 0.917 | 5 |
| c4_B - c7_B (most distant) | 0.848 | 7 |

Table S3. Mantel / Spearman statistics across 17 topologies. Only the true tree achieves a statistically significant Mantel p.

| Topology | Spearman ρ (Pearson sim) | Spearman ρ (cosine sim) | Mantel p |
|---|---|---|---|
| true_tree | +0.34 | +0.38 (p=0.04*) | 0.04* |
| alt_1-5 | -0.44 to +0.32 | -0.42 to +0.30 | all n.s. |
| fixed_1-5 | -0.08 to +0.26 | -0.08 to +0.25 | all n.s. |
| linear_1-5 | -0.28 to +0.32 | -0.27 to +0.34 | all n.s. |

Table S4. Expression recovery at optimal regularization strength (with-normal mode, all 8 clones and 320 replicates, star tree)

| Model | Best lambda | Mean Pearson r | 95% CI |
|---|---|---|---|
| Plain | 0.10 | 0.9419 | [0.928, 0.953] |
| Adaptive | 0.50 | 0.9403 | [0.926, 0.952] |
| Adaptive_v2 | 0.50 | 0.9403 | [0.926, 0.952] |
| Plain_debiased | 0.50 | 0.9369 | [0.922, 0.949] |
| Fused_ew | 0.01 | 0.9228 | [0.905, 0.938] |
| Elastic_net | 0.01 | 0.9336 | [0.918, 0.946] |
| Tree_delta | 0.05 | 0.9404 | [0.926, 0.952] |

Table S5. GSEA pathway recovery on synthetic edges (with-normal mode, pooled across 4 synthetic edges, 320 replicates, best lambda per model, star tree).

| Model | lambda | Sensitivity | F1 | MCC |
|---|---|---|---|---|
| Plain | 0.10 | 0.273 | 0.305 | 0.232 |
| Adaptive | 0.50 | 0.237 | 0.291 | 0.251 |
| Adaptive_v2 | 0.50 | 0.237 | 0.286 | 0.246 |
| Plain_debiased | 0.50 | 0.316 | 0.326 | 0.207 |
| Fused_ew | 0.01 | 0.273 | 0.299 | 0.207 |
| Elastic_net | 0.01 | 0.339 | 0.344 | 0.220 |
| Tree_delta | 0.05 | 0.283 | 0.321 | 0.233 |
| Baseline | — | 0.211 | 0.271 | 0.183 |

Table S6. Pathways altered for each synthetic edge.

| Synthetic clone | Parent | Assigned Hallmark sets |
|---|---|---|
| c7_A | c7 | HYPOXIA, GLYCOLYSIS, REACTIVE_OXYGEN_SPECIES_PATHWAY, FATTY_ACID_METABOLISM, OXIDATIVE_PHOSPHORYLATION |
| c7_B | c7_A | MYC_TARGETS_V1, G2M_CHECKPOINT, E2F_TARGETS, DNA_REPAIR, MYC_TARGETS_V2 |
| c4_A | c4 | TGF_BETA_SIGNALING, INFLAMMATORY_RESPONSE, COMPLEMENT, COAGULATION, IL6_JAK_STAT3_SIGNALING |
| c4_B | c4_A | INTERFERON_GAMMA_RESPONSE, TNFA_SIGNALING_VIA_NFKB, APOPTOSIS, PI3K_AKT_MTOR_SIGNALING, MTORC1_SIGNALING |
